# MITF, TFEB and TFE3 drive distinct adaptive gene expression programs and immune infiltration in melanoma

**DOI:** 10.1101/2024.12.23.629393

**Authors:** Diogo Dias, Erica Oliveira, Román Martí-Díaz, Sarah Andrews, Ana Chocarro-Calvo, Alice Bellini, Laura Mosteo, Yurena Vivas García, Jagat Chauhan, Linxin Li, José Manuel García-Martinez, José Neptuno Rodriguez-López, Silvya Stuchi Maria-Engler, Colin Kenny, Javier Martínez-Useros, Custodia García-Jiménez, Luis Sanchez-del-Campo, Pakavarin Louphrasitthiphol, Colin R Goding

## Abstract

Cells can contain multiple related transcription factors targeting the same sequences, leading to potential regulatory cooperativity, redundancy, competition or temporally regulated factor exchange. Yet the differential biological functions of co-targeting transcription factors are poorly understood. In melanoma, three highly related transcription factors are co-expressed: The mTORC1-regulated TFEB and TFE3, key effectors of a wide range of metabolic and microenvironmental cues assumed to perform similar functions; and MITF, that controls melanoma phenotypic identity. Here we reveal the functional specialization of MITF, TFE3 and TFEB and their impact on melanoma progression. Notably, although all bind the same sequences, each regulates different and frequently opposing gene expression programs to coordinate differentiation, metabolism, and protein synthesis, and qualitatively and quantitatively impact tumor immune infiltration. The results uncover a hierarchical cascade whereby microenvironmental stresses, including glucose limitation, lead MITF, TFEB and TFE3 to drive distinct biologically important transcription programs that underpin phenotypic transitions in cancer.

## Introduction

Co-expression of structurally related transcription factors in cells can lead to potential regulatory cooperativity, redundancy or competition, or sequential activity at the same regulatory elements^1^. This may allow flexibility in controlling common target genes, especially if the co-targeting transcription factors perform non-equivalent functions. However, our understanding of how functionally similar transcription factors perform distinct roles is limited. This is particularly important in cancer where disease progression is largely driven by microenvironment-mediated changes in cell phenotype^2-4^. For example, nutrient or oxygen limitation leads cells to employ pro-survival strategies including metabolic reprogramming and migrating to seek an improved environment^5,6^. Having multiple, highly related transcription factors targeting the same genes but eliciting different outcomes may be a key mechanism that allows cells to undergo appropriate non-genetic adaptations such as a switch from proliferation to invasion.

Melanoma represents an excellent model to understand the complex interplay between transcription factors, cell plasticity, the microenvironment and disease progression^7^. The microphthalmia-associated transcription factor MITF^8^, a lineage survival oncogene^9^, is a key determinant of phenotypic identity and plays a critical coordinating role in many aspects of melanoma biology^10^. MITF promotes survival^9,11^, proliferation^12,13^ and differentiation^14,15^, regulates metabolism^16-19^, controls DNA damage repair^20,21^, and suppresses senescence^22^ and invasion^23^. In vivo, MITF expression is dynamic^24^ and is suppressed by several microenvironmental stressses^10^ including nutrient limitation^25,26^, hypoxia^19,27^ and inflammatory signaling^25^ that slow proliferation but facilitate metastatic dissemination. However, while MITF expression is reduced in invasive cells^23^ that have elevated tumor-initiating ability^25,28^ and exhibit non-genetic therapy resistance^29-32^, for invasive cells to generate a new metastasis, MITF should be re-expressed to allow proliferation^15,33^. Despite these insights, our understanding of the molecular adaptations occurring as cells are subject to potentially lethal stresses during disease progression remains incomplete.

One possibility, yet to be fully explored, is that on downregulation of MITF, the MITF-related transcription factors TFEB and TFE3^10,34,35^ take on MITF functions. TFE3 and/or TFEB are key regulators of autophagy and lysosome biogenesis^34,36-38^, but are also implicated in many other cellular processes. For example, modulating the integrated stress response^39,40^, insulin expression^41^, the innate immune response^42,43^, recovery from exercise^44^ and lineage specification and stem cell fate^45,46^. Not surprisingly, given their importance, deregulation of TFEB and TFE3 is implicated in cancer^47-50,51^. Unlike the melanocyte/melanoma-specific isoform of MITF, in nutrient rich conditions TFE3 and TFEB are retained in the cytoplasm via 14-3-3 protein interaction^52^ following their phosphorylation by the Mammalian Target of Rapamycin Complex 1 (mTORC1)^34,53^. mTORC1-mediated phosphorylation of TFEB and TFE3 requires their recruitment to endolysosomal membranes by the RAGC/D GTPases^54^ where their phosphorylation is dependent on Folliculin (FLCN), a RAGC-activator^55^. On nutrient limitation, decreased mTORC1 activity and dephosphorylation^56,57^ permit TFEB and TFE3 to enter the nucleus to regulate transcription.

Here we investigate the role of TFEB and TFE3 in cells that also express MITF to understand better the potential of these related transcription factors to exhibit distinct and non-redundant roles in gene expression and melanoma progression.

## Results

### MITF promotes mTORC1 signaling

If starvation can trigger invasion, why would MITF^Low^ cells maintain invasiveness under nutrient-rich culture conditions? We hypothesised that MITF^Low^ cells might fail to sense or use nutrients to impact regulation of TFEB and TFE3. We therefore examined the effect of MITF depletion on mTORC1 that promotes protein translation by increasing S6 kinase-mediated activation of S6 and phosphorylation of 4EBP1^58^. Depletion of MITF in MITF^High^ IGR37 cells led to lower phosphorylation of S6 and 4EBP1 (Figure 1A). Consistent with previous results^59^, silencing MITF led to an increase in PTEN protein (Figure 1A, left) and mRNA (Figure 1A, right), an inhibitor of PI3 kinase upstream of mTORC1.

**Figure 1.**
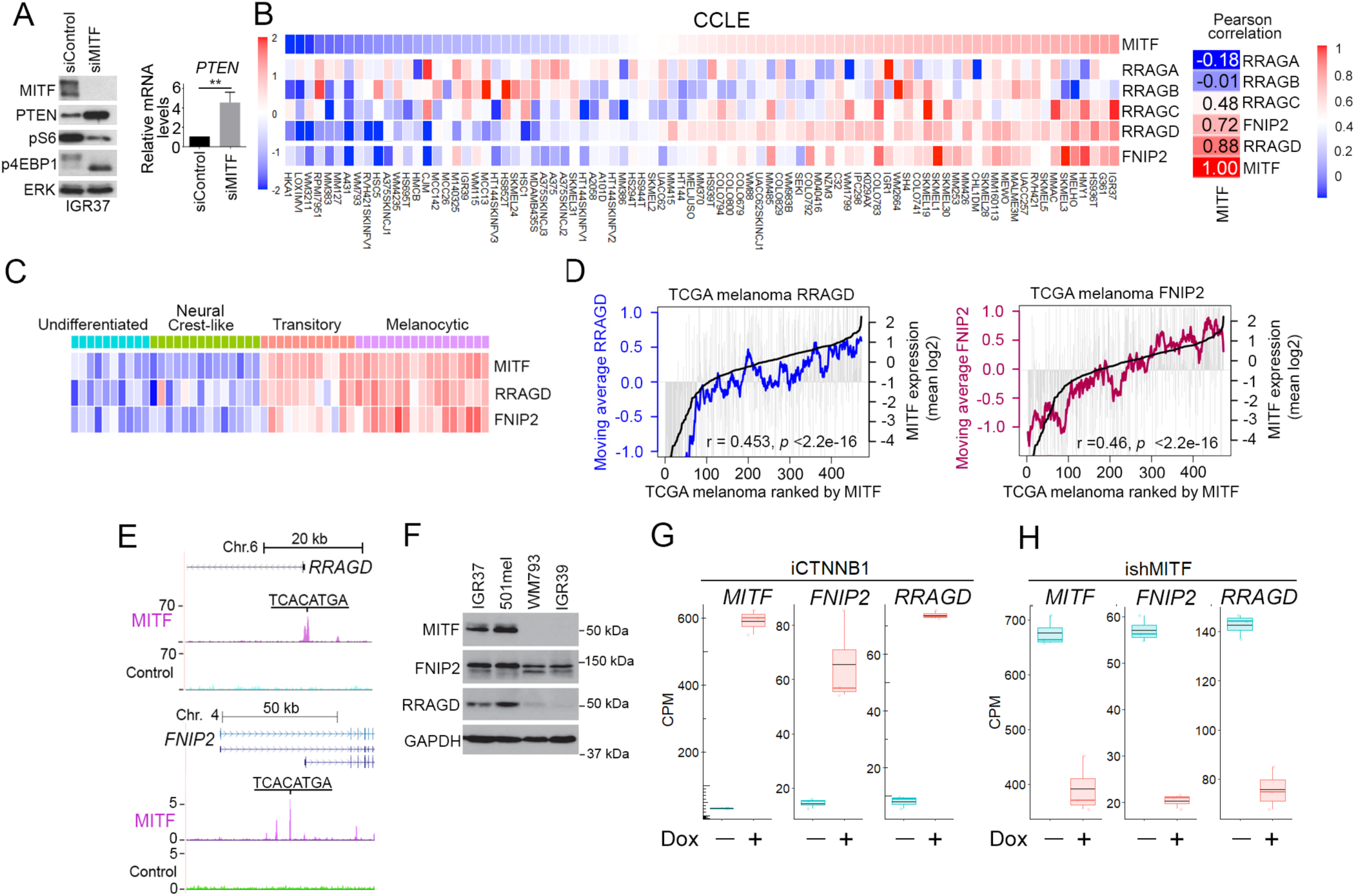
Control of mTORC1 by MITF. (A) Left, Western blot of IGR37 melanoma cell lines transfected with control or MITF-specific siRNA. Right, qRT PCR for *PTEN* mRNA from corresponding IGR37 cells. (B-C) Heatmaps showing relative mRNA expression in CCLE melanoma or Tsoi et al 53 melanoma cell lines. Figure 1B, right, Pearson correlation of genes expression. (D) TCGA melanomas ranked by MITF expression *MITF* (Black line). Grey bars indicate expression of *RRAGD* or *FNIP2* in each melanoma with the moving average of each per 20 melanoma window indicated by colored lines. (E) MITF ChIP-seq showing binding to the *RRAGD* or *FNIP2* genes. (F) Western blot of melanoma cell lines. (G, H) Box and whisker plots showing relative expression of *MITF*, *RRAGD* and *FNIP2* based on triplicate RNA-seq of IGR37 cells expressing doxycycline-inducible β-catenin (iCTNNB1) (G) or in MeWo cells after shRNA-mediated depletion of MITF (H). See also Figure S1.

To identify PTEN-independent mTORC1 inhibitory mechanisms, we examined the RAG GTPases and folliculin-interacting protein 2 (FNIP2) that modulate mTORC1 signaling at the lysosome^55,60-62^ to control TFEB and TFE3. In the Cancer Cell Line Encyclopedia (CCLE) expression of *FNIP2* and *RRAGD* and to some extent *RRAGC*, but not *RRAGA* or *RRAGB*, correlate with *MITF* expression (Figure 1B). The correlation was reproduced in the 53 melanoma cell lines described by Tsoi et al ^63^ that cluster into 4 distinct phenotypes (Figure 1C). *RRAGD* and *FNIP2* mRNA levels were strongly expressed in the MITF^High^ transitory and melanocytic phenotypes but not in the undifferentiated or neural crest-like phenotypes. The correlation between *MITF* and *RRAGD* and *FNIP2* mRNA expression was reproduced using The Cancer Genome Atlas (TCGA) melanoma cohort (Figure 1D). Importantly, ChIP-seq^64^ revealed MITF binding to the promoters of *RRAGD* and isoform 2 of *FNIP2* (Figure 1E) and *FNIP2*, and *RRAGD* expression were lower in the MITF^Low^ WM793 and IGR39 melanoma cell lines compared to MITF^High^ IGR37 and 501mel cells (Figure 1F). Doxycycline-mediated induction of β-catenin (CTNNB1), an MITF co-factor^65^ and activator of *MITF* expression^66^, in IGR37 cells induced *FNIP2* and *RRAGD* expression (Figure 1G), while depletion of MITF decreased their expression (Figure 1H). These data suggest that *FNIP2* and *RRAGD* are MITF target genes, and that signaling to TFEB and TFE3 would potentially be deregulated in MITF^Low^ melanoma cells.

### Differential regulation of TFEB and TFE3 by glucose limitation

Consistent with previous observations in melanoma^67^, in the TCGA melanoma cohort *TFE3* mRNA was inversely correlated with *MITF* (Figure 2A) while a moderate anti-correlation was also noted between *MITF* and *TFEB* (Figure 2B). By contrast, *TFEB* and *TFE3* mRNA correlated (Figure 2C). Using the Tsoi et al^63^ melanoma cell line panel (Figure 2D) *TFE3* mRNA was primarily expressed in the MITF^Low^ undifferentiated and neural crest-like phenotypes while *TFEB* mRNA expression was not phenotype restricted. In an additional panel of 12 cell lines, mRNA-seq indicated predominantly low *TFE3* expression in MITF^High^ cell lines, while *TFEB* mRNA expression was variable (Figure 2E). Western blotting (Figure 2F) revealed that TFEB protein was more abundant MITF-expressing cells, with SKMEL28 and SKMEL30 notable exceptions, but TFE3 expression was largely independent of MITF status. Thus, some cells express all three MITF family members, but in others only TFE3 is expressed. Any discordance between TFE3 or TFEB protein and mRNA expression may arise because unlike TFEB, under nutrient replete conditions TFE3 is subject to CUL^β−TrCP^-dependent ubiquitination and degradation^50^ and other factors such as differential translation efficiency may also play a role.

**Figure 2.**
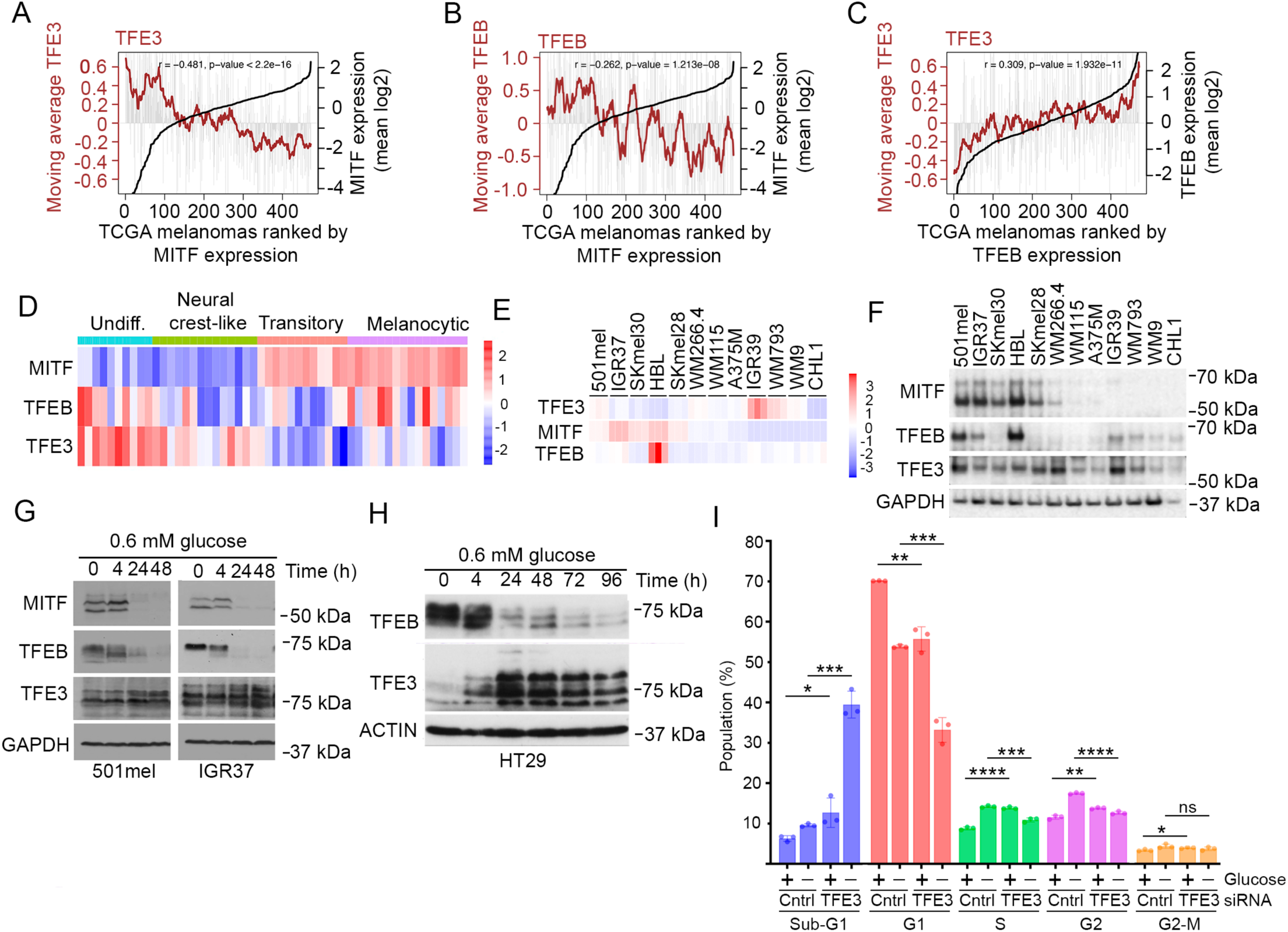
Differential expression of MITF, TFEB and TFE3 on glucose limitation. (A-C) TCGA melanomas ranked by expression of *MITF* (A,B) or TFEB (C) (Black lines). Grey Bars indicate expression of TFE3 (left and right) or TFEB (middle) in each melanoma with the moving average of each per 20 melanoma window indicated by colored lines. (D-E) Heatmaps showing relative gene expression in the Tsoi et al melanoma cell line panel (D) or an inhouse panel (E). (F) Western blot of panel of melanoma cell lines. (G-H) Western blot of 501mel melanoma (G) or HT29 colorectal cancer cells (H) grown in low glucose medium. (I) Flow cytometry of IGR37 cells transfected with control siRNA or a siTFE3 pool for 48 h before being placed in glucose-free DMEM for 6 h. Error bars indicate S.D. N=3. *= *p* <0.05; **= *p* <0.01; ***= *p* <0.001; ****= p <0.0001 *ns* = not significant, Student t-test.

Immunofluorescence using the MITF^High^ (501mel and IGR37) and MITF^Low^ (IGR39 and WM793) cell lines revealed low nuclear expression of TFEB (Supplemental Figure 1A) which was increased using Torin, an mTOR inhibitor. Similarly, TFE3 was also largely nuclear in the MITF^High^ 501mel and IGR37 cells with expression being increased by Torin. However, unlike TFEB, in IGR39 cells TFE3 was predominantly cytoplasmic but accumulated in the nucleus following Torin treatment. In WM793 cells the low levels of cytoplasmic TFE3 exhibited a moderate nuclear accumulation in response to Torin.

Western blotting of fractionated extracts from 501mel cells revealed that Torin induced a loss of the slow-migrating phosphorylated form of TFEB and its nuclear accumulation (Supplemental Figure S1B). On depletion of MITF there was a similar accumulation of nuclear TFEB using Torin, but cytoplasmic phosphorylated TFEB levels were reduced. Notably, immunohistochemistry using tissue sections from three different human melanomas (Melanomas 1-3) revealed a significant proportion of melanoma cells exhibiting nuclear staining of TFEB (Supplemental Figure S1C) suggesting that in vivo mTORC1 signaling is suppressed. This is consistent with 63% of melanoma metastases exhibiting nuclear TFE3 staining^68^. While the high nuclear MITF in Melanoma 2 correlated with the lowest total TFEB staining, as the same sections were not probed with both antibodies the relationship between TFEB and MITF expression and subcellular localization in individual cells in vivo remains uncertain.

Glucose limitation, frequently found in tumors, regulates TFEB nuclear/cytoplasmic shuttling^69^. Consistent with previous work^26^, low glucose, silenced MITF (Figure 2G), and TFEB exhibited at 4 h a faster mobility characteristic of its nuclear accumulation^55,69,70^. However, by 24 h TFEB expression was strongly suppressed (Figure 2G). By contrast, TFE3 expression was not silenced and after 48 h in low glucose it was the only MITF-family member expressed. Similar results were obtained in HT29 colon cancer cells (Figure 2H) where glucose depletion downregulated TFEB but upregulated TFE3. We therefore asked whether depletion of TFE3 might affect the cell cycle or survival in glucose deprived cells. The results (Figure 2I) revealed that in low glucose, depletion of TFE3 led to a dramatic increase in the sub-G1 population and corresponding decrease in G1 cells, suggesting that TFE3 plays a key pro-survival role.

### MITF, TFE3 and TFEB bind the same genes

Under glucose limitation, MITF and TFEB expression is suppressed, but TFE3 is maintained. Although all three proteins have an identical basic region that makes sequence specific DNA contacts (Figure 3A), differential interaction with cofactors might enable them to regulate differentially subsets of genes. We therefore used 501mel human melanoma cells, in which we previously mapped genome-wide binding of MITF ^64,71^, to express doxycycline (Dox)-inducible HA-tagged TFEB (Figure 3B) and performed ChIP-seq using the highly specific HA antibody. Since mTORC1 inhibits TFEB we performed ChIP-seq using 0 ng or 5 ng of Dox +/- 250 nM Torin that induced nuclear accumulation of HA-tagged TFEB (Supplemental Figure S2A). Replicate ChIPs showed a good concordance (Supplemental Figure S2B). The results (Figure 3C) revealed substantial genome occupancy without Dox and Torin consistent with TFEB cycling in and out of the nucleus under nutrient rich conditions ^69,72^. Nevertheless, Torin treatment increased the mean peak score and the number of binding sites recognised (Supplemental Figure S2C). Using 5 ng Dox, HA-TFEB DNA binding increased to 9576 sites (Supplemental Figure S2D) with a higher mean peak score (Figure 3C), consistent with previous work using inducible MITF^64,71^. Using both Dox and Torin led to 39,582 sites being occupied (Supplemental Figure S2D) of which 9522 were bound in common with the Dox-only condition. These data were reflected in the read density maps (Supplemental Figure S2E).

**Figure 3.**
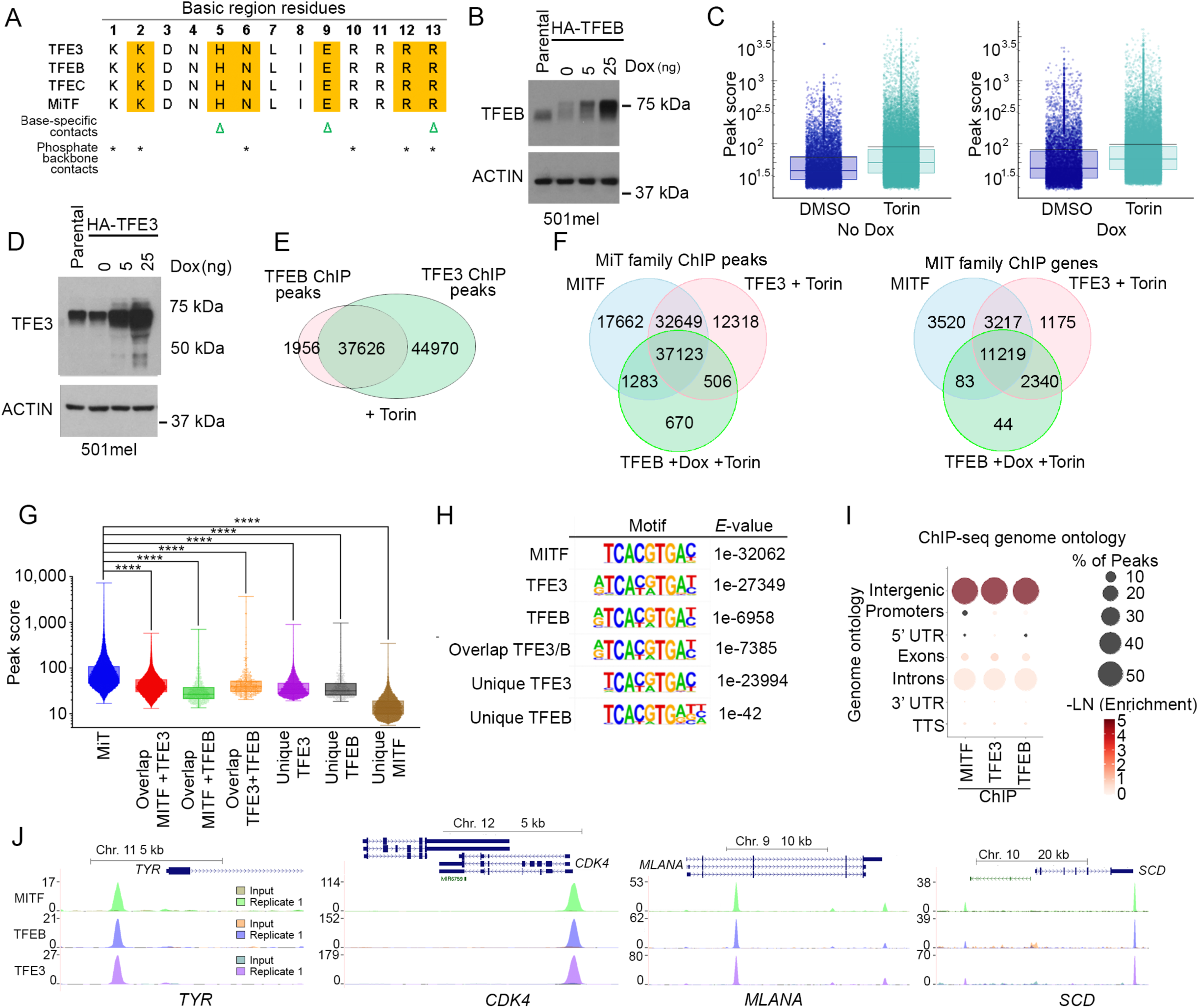
MITF family members bind a similar repertoire of genes. (A) Sequences of MITF family member basic regions. (B) Western blot showing expression of doxycycline-inducible HA-tagged TFEB in 501mel cells. (C) Jitter plots showing peak scores of HA-TFEB ChIP-seq in 501mel cells +/- 250 nM Torin for 6 h and either 0 ng or 5 ng doxycycline. Black lines indicate mean. Boxes indicate interquartile range and median. (D) Western blot showing expression doxycycline-inducible HA-tagged TFE3 in 501mel cells. (E-F) Venn diagrams showing the number of statistically significant ChIP peaks or genes bound by MITF family members. HA-TFEB and HA-TFE3 ChIPs were performed in the presence of 250 nM Torin. No doxycycline was used for the HA-TFE3 ChIP and 5 ng for the TFEB ChIP. The MITF data was taken from Louphrasitthiphol et al, 2020. (G) Relative peak scores at sites bound by each factor alone or indicted combinations. Boxes indicate interquartile range and median. **** indicates *p-*value < 0.001 One-way ANOVA. (H) Motifs identified beneath ChIP peaks for MITF, TFEB and TFE3. (I) Genome ontology of binding sites identified for each factor. (J) Color coded ChIP profiles for input control and HA-TFEB, HA-TFE3 and HA-MITF. Only one replicate is shown. Similar results were obtained for replicate 2. See also Figure S2.

We next repeated the ChIP-seq using HA-TFE3 in the absence of Dox to maintain TFE3 at levels seen in parental cells (Figure 3D). Torin increased nuclear HA-TFE3 (Supplemental Figure S2F) with good concordance between replicates (Supplemental Figure S2G, H). We then compared the results with those of TFEB (Figure 3E), and also undertook a 3-way comparison with a previously published^64^ MITF ChIP (Figure 3F). These analyses revealed 37,123 sites (Figure 3F, left) at 11,219 genes (Figure 3F, right) bound by all three factors, but also many peaks apparently bound by each alone or by any two factors. For example, MITF and TFE3 share 32,649 peaks at 3217 genes that are not called in the TFEB ChIP-seq.

We noted that the peak score of sites bound by all three factors was higher than those bound by any single or combination of factors (Figure 3G). Thus, rather than being biologically relevant, peaks unique to each factor were more likely ‘bioinformatically unique’. Examination of binding at many loci, including those in Figure S2I, supported this notion. For example, at the *ATAD3B* gene (Figure S2I, left) an ‘MITF unique’ peak is also present in the TFE3 ChIP but was not called because of elevated reads in the TFE3 input control. It was not called in the TFEB ChIP owing to reduced ChIP efficiency and higher reads in the input control. Similarly, a ‘TFE3 unique’ peak upstream from the *BLOC1S6* gene (Figure S2I, right), is present in the MITF ChIP but is not called owing high input control reads. At both loci, only a partial consensus MITF family binding site was found beneath the ‘unique’ peaks. Using an alternative differential tag-density approach using the ChIP-seq data from the other two factors as an input we did not detect peaks unique to any one factor (Figure S2J). Overall, while we cannot rule out binding by individual factors to a few unique genes, most potential regulatory sites are likely bound by all three factors.

All three factors recognised a similar 8 bp consensus TCACGTGA (Figure 3H) previously identified for MITF^73^, also known as the CLEAR box^36^, and exhibited a similar genomic distribution (Figure 3I). As examples, binding to four known MITF target genes is shown in (Figure 3J).

To ask whether factor exchange occurs, we interrogated a recently published CUT&RUN data set^74^ obtained for MITF and TFE3 in SKMEL28 WT cells or a previously published^75^ *MITF* KO. SKMEL28 cells do not express TFEB (Figure 2F), which simplifies data interpretation. The tag density heatmaps in Figure S2K reveal the *MITF* KO exhibits greatly reduced MITF binding with residual binding reflecting some antibody cross-reactivity with another MITF family member, as reported^74^. Importantly, the low TFE3 binding in the WT cells increased substantially in the *MITF* KO, likely owing to absence of competition and because low MITF reduces mTORC1 activity (Figure 1) leading to elevated nuclear accumulation of TFE3. As examples, at *DCT* (Supplemental Figure S2L) or *TRPM1* (Supplemental Figure S2M) there is reduced MITF binding in the *MITF* KO, but increased TFE3 occupancy. By contrast, at the *GPR143* locus, TFE3 binding occurs neither in the WT nor *MITF* KO cells (Supplemental Figure S2N). It is possible that loss of MITF leads to nucleosome remodeling rendering sites at genes like *GPR143* inaccessible to TFE3.

### MITF family members regulate different gene sets

Although MITF, TFEB, and TFE3 bind similar targets, DNA binding does not necessarily equate to regulation^76,77^. We therefore examined the contribution of each factor to gene expression using MITF^High^ 501mel human melanoma cells that express all three factors (Figure 2F). We initially performed RNA-seq on 501mel cells exposed to HEPES-buffered saline solution (HBSS) that contains glucose but lacks amino acids. The results revealed over 2,400 differentially expressed genes DEGs (fold change (FC) ≥ 2; *p* <0.05) (Figure 4A, B; Supplemental Table S1). Gene Set Enrichment Analysis (GSEA) (Figure 4C, D) revealed downregulation of the HALLMARK E2F1 and MYC TARGET gene sets, as well as DNA repair genes previously reported to be targets of MITF^20,22,78,79^.

**Figure 4.**
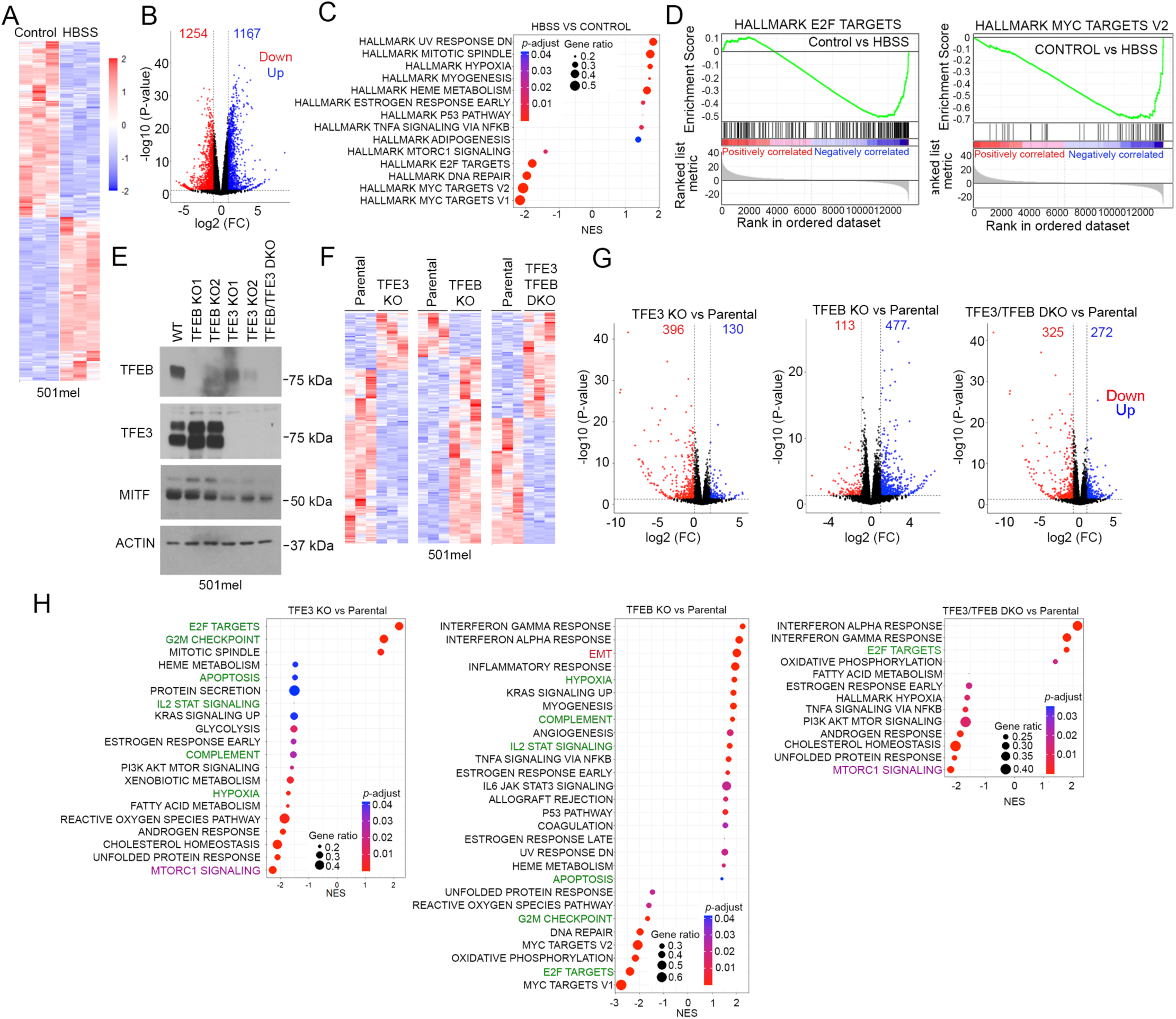
Differential gene expression in *TFEB*, *TFE3* and *TFEB/TFE3* KO human melanoma cells. (A) Triplicate RNA-seq showing differential gene expression of control 501mel cells or cells grown in HBSS for 12 h. (B) Volcano plot showing numbers of significantly (FC ≥ 2, *p* = <0.05) differentially expressed genes comparing control and HBSS-treated cells. (C) GSEA plots showing significantly differentially enriched gene sets comparing control and HBSS-treated 501mel cells. (D) GSEA plots comparing control versus HBSS-treated 501mel cells. (E) Western blot using parental and *TFEB* and/or *TFE3* 501mel KO cell lines. (F) Triplicate RNA-seq showing differential gene expression (FC ≥ 2, *p* = <0.05) of parental or *TFEB* and/or *TFE3* KO 501mel cells. (G) Volcano plots showing numbers of significantly (FC ≥ 2, *p* = <0.05) differentially expressed genes comparing parental cells and indicated 501mel KOs. (H) GSEA plots comparing parental or indicated *TFEB* and/or *TFE3* 501mel KO cells. See also Figure S3 and Table S1.

We next used CRISPR/Cas9 to knockout (KO) each gene and were successful in inactivating *TFEB* and *TFE3*, and in making a *TFE3/TFEB* double knock-out (DKO) (Figure 4E) but were unable to generate an *MITF* KO. The *TFEB* KO cell lines exhibited modest upregulation of TFE3 but no change on MITF. The *TFE3* KO cells moderately downregulate TFEB, especially clone 2, with some suppression of MITF. RNA-seq of *TFEB* (clone 2) and *TFE3* (clone 1) KOs, and the DKO grown under standard culture conditions revealed changes in gene expression (Figure 4F, G) consistent with TFEB and TFE3 possessing transcriptional activity under nutrient replete conditions. Importantly, GSEA (Figure 4H) revealed that KO of *TFE3* or *TFEB* had different effects on gene expression. The *TFE3* KO exhibited enrichment of the E2F TARGETS and G2M CHECKPOINT gene sets, and downregulation of gene sets including HYPOXIA, COMPLEMENT, IL2/STAT SIGNALING and APOPTOSIS (Figure 4H, highlighted in green). By contrast, in the *TFEB* KO, these gene sets were regulated in the opposite way. In addition, MTORC1 SIGNALING was downregulated in the *TFE3* KO and the *TFE3/TFEB* DKO, but not the *TFEB* KO. Many gene sets displaying altered expression in the single KOs were not differentially expressed in the DKO, suggesting the effect of one KO was partly reversed by KO of the related gene.

Torin treatment of the parental cells or the three KO cell lines led to altered gene expression (Supplemental Figure S3A, B) and as expected in the parental cells GSEA revealed significant downregulation by Torin of the mTORC1 SIGNALING, MYC and E2F TARGET gene sets (Supplemental Figure S3C) and a moderate enrichment of the EMT gene set.

Remarkably, all differentially expressed gene sets were downregulated in Torin-treated *TFE3* KO cells, and none were significantly upregulated (Supplemental Figure S3D). The *TFE3* KO therefore appears to amplify the effect of Torin, leading to suppression of gene sets associated with mTORC1 SIGNALING and CHOLESTEROL HOMEOSTASIS, as well as E2F and MYC TARGETS. The enrichment of the EMT gene set in parental cells exposed to Torin was reversed in the Torin-treated *TFE3* KO.

Using the *TFEB* KO cells (Supplemental Figure S3E), we again obtained a result largely the opposite to that seen using the *TFE3* KO. For example, enrichment of the MTORC1 SIGNALING gene set was reduced in the parental and *TFE3* KO cells by Torin but was increased in *TFEB* KO cells. By contrast, the MYC TARGETS V1 gene set was decreased in the parental, *TFE3*, *TFEB* KO and DKO cell lines on Torin treatment (Supplemental Figure S3F).

Collectively these data indicate that while both TFE3 and TFEB bind a similar repertoire of target genes their effects on gene expression may be different. However, data interpretation is complicated by 501mel cells expressing MITF. We therefore used a previously described^80^ murine B16-F10 cell line bearing a frameshift mutation in the *MITF* gene (Figure 5A). Note that close inspection reveals a minor fast-migrating band cross-reacting with the anti-MITF antibody in the *MITF* KO, suggesting they may express a low level of a truncated form of MITF as has been observed previously^75^ in an SKMEL28 *MITF* KO cell line. We nevertheless used CRISPR/Cas9 to KO *TFE3* and *TFEB* in the parental and *MITF* KO B16-F10 cells (Figure 5B).

**Figure 5.**
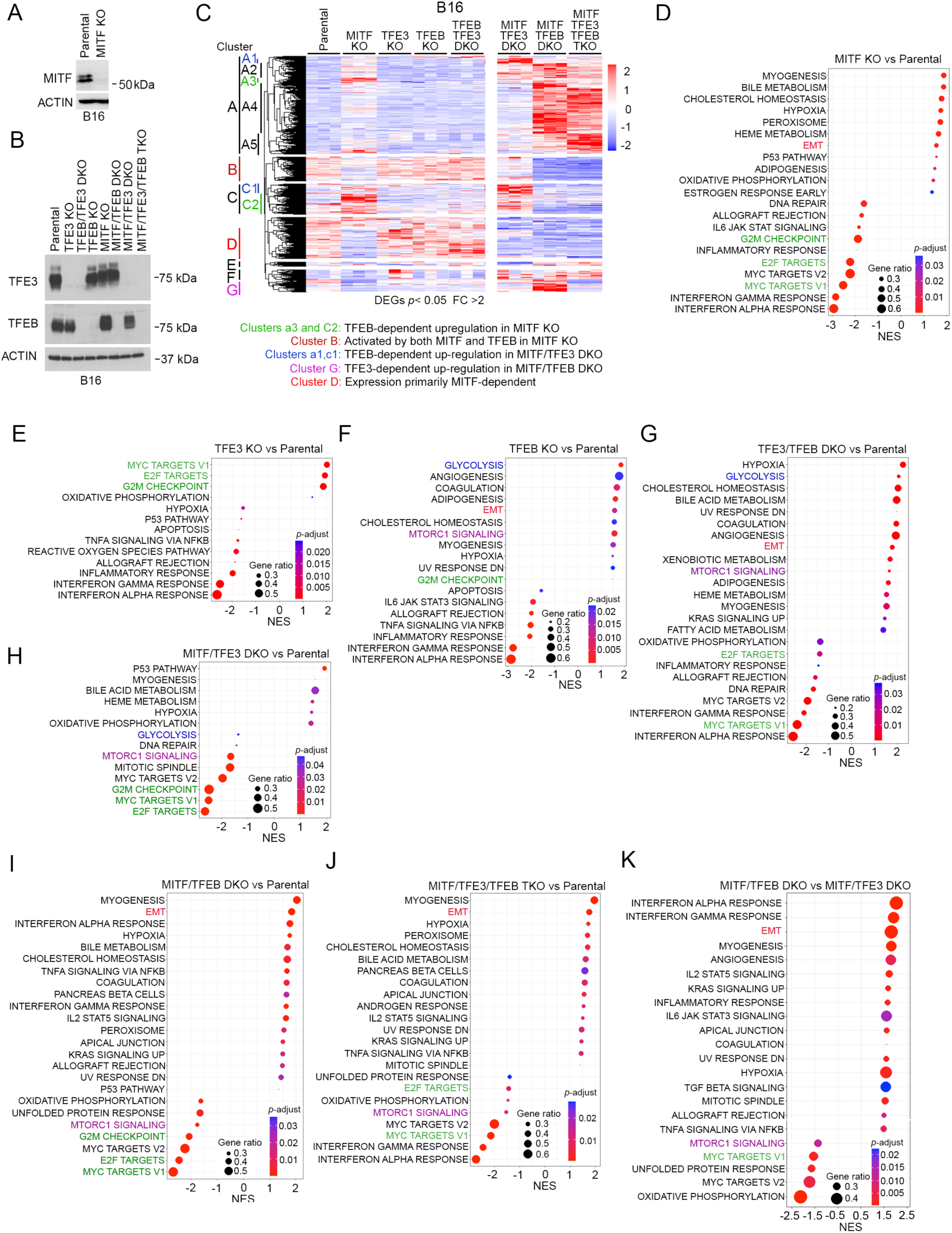
TFE3 and TFEB are not equivalent in gene regulation. (A) Western blot using B16-F10 parental and functional *MITF* KO cell line. (B) Western blot using parental and indicated B16-F10 KO cell lines. (C) Triplicate RNA-seq showing differential gene expression between parental and indicated B16-F10 KO cell lines. (D-K) GSEA plots showing significantly differentially enriched gene sets comparing parental or indicated *MITF*, *TFEB* and *TFE3* B16-F10 KO cell lines. See also Figures S4 and S5, and Table S2.

RNA-seq of each cell line grown under standard culture conditions revealed robust remodeling of the transcriptional landscape (Supplemental Figure S4A-G). Notably, compared to the *MITF* KO (Supplemental Figure S4A), fewer genes were differentially expressed in the *TFE3* and *TFEB* single KOs (Supplemental Figure S4B,C). The *MITF/TFEB* DKO (Supplemental Figure S4F) or the TKO (Supplemental Figure S4G) also exhibited more differentially expressed genes, suggesting that some genes were regulated by more than one family member. Hierarchical clustering of the RNA-seq of the parental and KO cells revealed a complex gene expression pattern (Figure 5C) with differential expression of genes in each cluster, labelled A-G, provided in Supplementary Table S2. For simplification, we only discuss a subset of the more interesting features.

First, KO of *MITF* has a bigger impact than KO of either *TFEB* or *TFE3* alone. For example, genes within clusters A3 and C2 are more upregulated in the *MITF* KO than in the *TFE3* or *TFEB* KOs (Figure 5C, green). Similarly, many genes downregulated in the *MITF* KO in clusters B and D are unaffected or increased in the *TFEB* or *TFE3* single KOs. The limited effect of the *TFE3* and *TFEB* single KOs may reflect that gene expression was measured in cells in nutrient rich medium where the TFEB/TFE3 inhibitor mTORC1 is active. Second, the upregulation of MITF-repressed genes in clusters A3 and C2 is reversed in the *MITF*/*TFEB* DKO or the TKO, indicating that TFEB upregulates these genes in the absence of *MITF*. Third, cluster B (Figure 5C, brown) contains genes moderately downregulated in the *MITF* KO but strongly downregulated in the *MITF/TFEB* DKO and TKO. Thus, MITF, and TFEB in the absence of MITF, positively regulate this gene set. Fourth, genes within clusters A1 and C1 (Figure 5C, blue) were upregulated in the *MITF*/*TFE3* DKO compared to the *MITF* KO, with the effect of the DKO being reversed in the TKO. MITF and TFE3 therefore prevent upregulation of these genes by TFEB. Fifth, cluster G genes (Figure 5C, pink) were upregulated in the *MITF/TFEB* DKO but not in the TKO. These are therefore controlled by TFE3 in the absence of both MITF and TFEB, as occurs when on glucose depletion. Finally, Cluster D genes (Figure 5C, red) downregulated in the *MITF* KO are largely unchanged in the *TFEB, TFE3* or *TFEB*/*TFE3* KOs. These genes appear to be MITF-dependent, but independent of TFEB and TFE3. Overall, each member of the MITF transcription factor family may compensate for another at some genes, but elsewhere they may play opposing roles.

GSEA analysis revealed MYC TARGETS, E2F TARGETS and G2M CHECKPOINT gene sets downregulation in the *MITF* KO (Figure 5D), but enrichment in the *TFE3* KO (Figure 5E), indicating opposing regulation of these pro-proliferative gene sets by MITF and TFE3. The MYC and E2F TARGETS gene sets were unchanged in the *TFEB* KO (Figure 5F).

The MTORC1 SIGNALING gene set was enriched in the *TFEB* KO and *TFE3*/*TFEB* DKO (Figure 5F) but suppressed in the *MITF*/*TFEB* DKO (Figure 5I), the *MITF*/*TFE3* DKO (Figure 5H) and the TKO (Figure 5J). This indicates the enrichment in the *TFEB* KO was dependent on MITF and to some extent on TFE3.

The GLYCOLYSIS gene set was enriched in the *TFEB* KO (Figure 5F) and *TFEB*/*TFE3* DKO (Figure 5G), but not in the TKO. No enrichment was observed in the *MITF*/*TFEB* DKO (Figure 5I), and GLYCOLYSIS was modestly downregulated in the *MITF*/*TFE3* DKO (Figure 5H). These data suggest TFEB restricts MITF-driven activation of glycolysis-related genes.

Finally, the INTERFERON ALPHA and GAMMA RESPONSE gene sets were downregulated in all single KOs, the *TFE3*/*TFEB* DKO (Figure 5G) and the TKO (Figure 5J). Comparing the *MITF*/*TFEB* DKO (Figure 5I) to the TKO (Figure 5J) suggests that in the absence of MITF and TFEB, TFE3 controls the INTERFERON RESPONSE gene sets.

A key conclusion is that TFE3 and TFEB behave very differently, a conclusion supported by comparing the genes significantly differentially regulated by the *TFE3* KO or *TFEB* KO in the *MITF* KO background (Supplemental Figure S4H). GSEA analysis (Figure 5K) revealed enrichment of the EMT, HYPOXIA and INTERFERON ALPHA and GAMMA RESPONSE gene sets and robust downregulation of mTORC1 SIGNALING, MYC TARGETS, UNFOLDED PROTEIN RESPONSE and OXIDATIVE PHOSPHORYLATION.

CyQuant assays revealed proliferation was reduced in the *MITF* KO, increased in the *TFE3* KO and unaffected in the *TFEB* KO or *TFE3/TFEB* DKO (Supplemental Figure S4J). By contrast the *MITF/TFE3* and *MITF/TFEB* DKOs and the TKO exhibited severely reduced proliferation. However, to establish cell lines, cells must proliferate. Consequently, any effect of each KO on proliferation may be underestimated.

The genes differentially regulated by combinations of MITF family members includes both those directly bound by MITF, TFEB and TFE3 and those where no binding was observed. However, regulation of genes by one factor but not another could arise on genes differentially bound by the transcription factors, and not because they have fundamentally different roles in regulating genes commonly bound by all three. To address this issue, we analysed the differential expression of genes in each KO cell line using only the 11,219 genes bound by all three factors (Figure 3F). The results, show major differences in gene expression that fall broadly into 10 clusters (Supplemental Figure S5A; Supplemental Table S3). For example, cluster 7, enriched in genes implicated in EMT and TGFβ signalling (Supplemental Figure S5B), is strongly upregulated in the *MITF* KO, but not in the *MITF*/*TFEB* DKO or the TKO (Supplemental Figure S5A). Similarly, genes in cluster 8, enriched in TGFβ signaling, are downregulated only in the *MITF*/*TFE3* DKO, whereas cluster 1 genes, enriched in mTORC1 and TNFα signalling, are only downregulated in the *MITF*/*TFEB* DKO and the TKO. The results strongly support the conclusion that genes bound by each MITF-family member may nevertheless be differentially regulated by them alone or in combination.

### MITF family genes differentially impact metabolism

The GSEA analysis indicated that MITF controls glycolysis-related gene expression. We therefore undertook Seahorse assays on the parental and MITF-family KO cell lines to measure the extracellular acidification rate (ECAR) and oxygen consumption rate (OCR), indirect measures of glycolysis and oxidative phosphorylation respectively. The ECAR (Figure 6A, left panel; Figure 6B) was lowest in the parental cells and moderately increased in all KO cell lines, and especially in the *TFE3* KO and *MITF*/*TFEB* DKO, with the TKO exhibiting the least increase. Treatment with oligomycin, an inhibitor of mitochondrial reversible ATP synthase that uncouples oxidative phosphorylation, revealed that all cells retained a glycolytic reserve proportional to basal glycolysis. Treatment with rotenone and antimycin A to block the electron transport chain had a moderate effect only in the *TFEB*/*TFE3*, *MITF*/*TFE3* and TKO cell lines. Measurement of the OCR (Figure 6A, right panel; Figure 6C) revealed greater differences between cell lines. Whereas parental B16-F10 cells have a low basal OCR, a striking increase was detected in the *MITF* KO and to some extent in the *MITF*/*TFE3* and *MITF*/*TFEB* DKOs. The TKO exhibited a reduced increase compared to the *MITF* KO or *MITF*/*TFEB* and *MITF*/*TFE3* DKOs. Thus, while MITF suppresses oxidative phosphorylation it is promoted by TFEB and TFE3 in the absence of MITF. The minor increase in basal OCR in the *TFE3* and *TFEB* KOs was not significant compared to the TKO. Consistent with these data, the *MITF* KO produced 60% more ATP than the parental cells, with the most significant increase arising from mitochondrial ATP production (Figure 6D). For the remaining KO cell lines, the ratio of mitochondrial to glycolytic ATP production was similar to the parental cells. However, the increased mitochondrial ATP production by the *MITF* KO cells was reduced in the TKO, indicating that much of the increase in the MITF KO depends on TFEB and TFE3. In agreement, many genes in the HALLMARK OXIDATIVE PHOSPHORYLATION gene set were upregulated in the *MITF* KO, but not in the *MITF*/*TFEB* DKO or the TKO (Figure 6E). Increased expression of these genes in the *MITF* KO is therefore dependent on *TFEB*.

**Figure 6.**
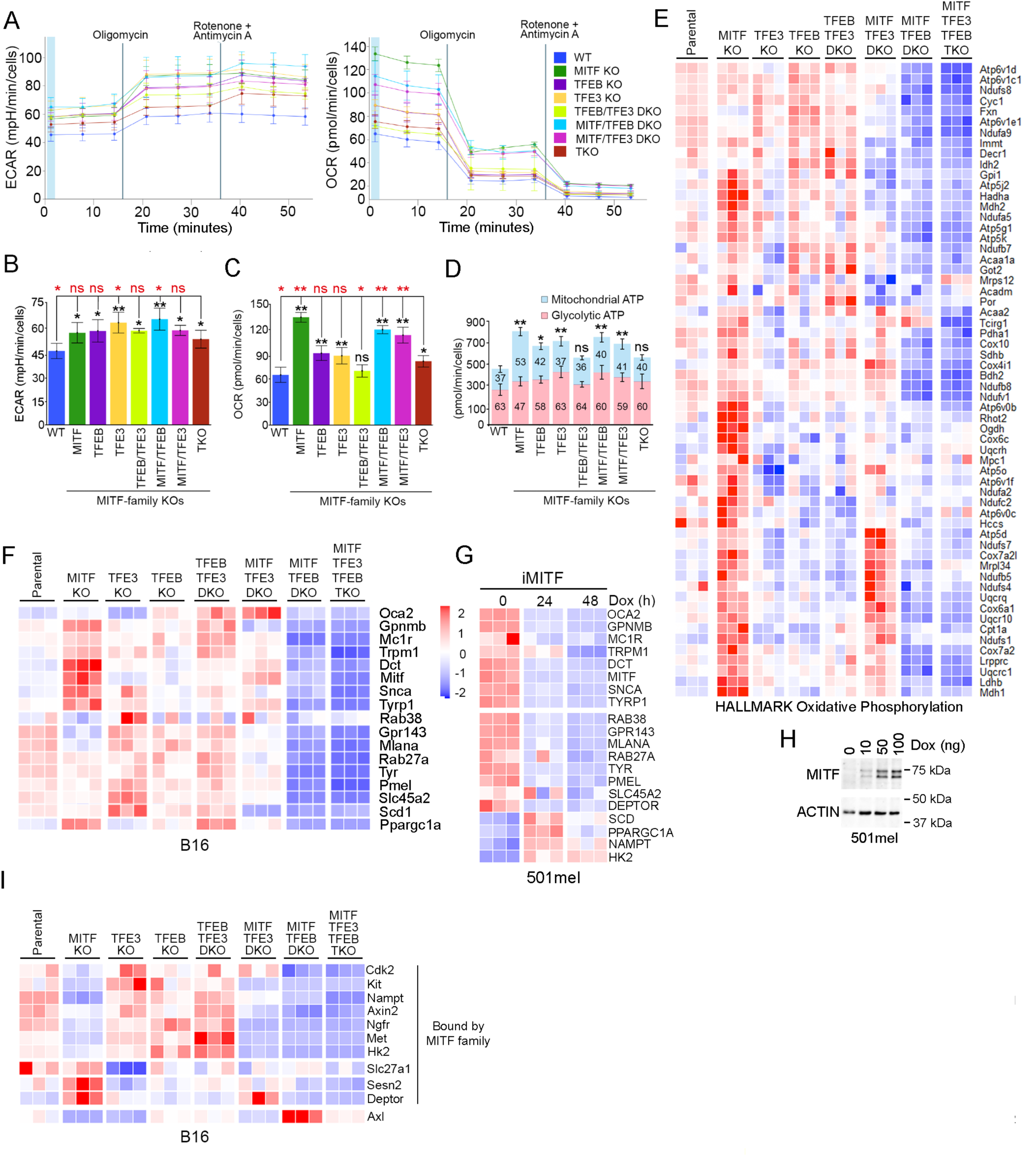
Differential regulation of metabolism and differentiation by MITF, TFEB and TFE3. (A) Seahorse assays measuring ECAR and OCR in B16-F10 WT or KO cell lines. Error bars = SD; n=3. (B, C) Seahorse assays summarising differential basal ECAR (B) and OCR (C) between B16-F10 WT or KO cell lines. Error bars =SD; n=3; ns = non-significant, *= p<0.05, **= p<0.01 (black, comparison with parental cells; red, comparison with TKO); Student t-test. (D) ATP production from glycolysis or oxidative phosphorylation by indicated cell lines. Numbers indicate % derived from each source. Error bars =SD; N=3; ns = non-significant, *= p<0.05, **= p<0.01, Student t-test comparing to parental WT B16-F10 cells. (E-G) Triplicate RNA-seq showing relative gene expression in parental and B16-F10 KO cell lines of a subset from the HALLMARK OXIDATIVE PHOSPHORYLATION gene set (E) or indicated genes (F), or 501mel cells in which MITF has been induced using 100 ng doxycycline (G). (H) Western blot of 501mel cells expressing ectopic HA-tagged MITF induced with 100 ng doxycycline for 16 h. (I) Triplicate RNA-seq showing relative gene expression parental and B16-F10 KO cell lines. See also Figure S6.

### Differential regulation of key target genes by the MITF-family

Multiple promoters drive expression of distinct MITF isoforms in different cell types^10^. All three factors bind the *MITF* gene (Supplemental Figure S6A) at consensus sequences (TCACATGA or ACACGTGA) upstream of the melanocyte specific MITF-M isoform. Inactivation of *MITF* increased *MITF* mRNA levels (Supplemental Figure S6B), consistent with previous work^19^, as well as expression of *TFEB* and *TFE3*. The increased *MITF* mRNA in the *MITF* KO was abrogated in the *MITF/TFEB* DKO and further reduced in the TKO (Supplemental Figure S6B). TFEB and TFE3 therefore positively regulate *MITF* expression in the absence of MITF. No binding by any MITF family member was detected at the *TFEB* and *TFE3* genes (Supplemental Figure S6C,D) and only one weak non-consensus binding site around 18 kb downstream from *TFEB* was identified. Any change in *TFEB* or *TFE3* expression in the KOs is therefore likely indirect.

We next examined a set of genes relevant to melanocyte/melanoma biology known to be directly bound and positively regulated by MITF (Figure 6F) including *Ppargc1a*^16,17^ and *Scd1*^18^. The results were surprising. In the *MITF* KO some genes implicated in melanocyte differentiation, such as *Mc1r*, *Dct*, *Tyrp1*, *Trpm1*, were upregulated, while others such as *Tyr*, *Pmel*, and *Slc45a* were largely unchanged. Only *Mlana*, *Gpr143* and *Rab38a* were moderately downregulated. This suggests that in proliferating B16-F10 cells MITF is a negative regulator at some differentiation genes. This interpretation was confirmed using human 501mel melanoma cells expressing Dox-inducible murine MITF (Figure 6G, H) where induction of MITF suppressed all differentiation-associated genes. By contrast, pro-proliferation genes such as *SCD*, *PPARGC1A*, *NAMPT* and *HK2* were upregulated. Although many differentiation genes were upregulated in the *MITF* KO, expression these genes, including *Scd1* and *Ppargc1a,* were downregulated in the *MITF*/*TFEB* DKO and the TKO (Figure 6F). Thus, in the absence of MITF, basal expression of these genes, which are all bound by all three MITF family members, was dependent on TFEB. Example binding profiles for *TYR*, *MLANA* and *SCD* are shown in Figure 3J, and for *PMEL* in Supplementary Figure 6E.

That MITF is pro-proliferative in B16-F10 cells was reinforced by examining a broader set of proliferation-associated targets including *Cdk2*^13^ and *Met*^81^, the receptor tyrosine kinases *Kit* and *Ngfr*, and *Nampt*, *Axin2* and *Hk2*, all of which are bound by all three family members. All were moderately downregulated in the *MITF* KO (Figure 6I), as well as in the *MITF/TFE3* and *MITF/TFEB* DKOs, and the TKO. These results together indicate that at some differentiation genes MITF may play a repressive role in proliferating melanoma cells, while promoting expression of pro-proliferation genes.

MITF^High^ cells uptake long-chain fatty acids via fatty acid transporter proteins (FATPs) to fuel melanoma progression^82-84^. Inactivation of *MITF* or *TFEB* had little effect on expression of *Slc27a1* (Figure 6I), that encodes FATP1 and is bound by all MITF family members (Supplementary Figure S6E). However, *Slc27a1* expression is strongly downregulated in the *TFE3* KO (Figure 6H).

Loss of *MITF* also upregulated two mTORC1 inhibitors, *Sesn2*^85-88^ and *Deptor*^89^, consistent with MITF promoting mTORC1 signaling (Figure 1). Both genes are bound by all family members at consensus TCACGTGG (*Sesn2*) or TCATGTGA (*Deptor*) motifs (Supplementary Figure S6E). Regulation of *DEPTOR* by MITF was confirmed in IGR37 cells by qRT PCR following depletion of MITF (Supplementary Figure S6F) and by Western blotting (Supplementary Figure S6F, lower panel). The elevated *Sesn2* and *Deptor* expression observed in the *MITF* KO was not seen in the TKO, implicating other family members in in their upregulation.

Interestingly, *AXL*, encoding a key receptor tyrosine kinase linked to therapy resistance^29,30^, is expressed in MITF^Low^ melanoma cells and can be activated by unsaturated fatty acids^83^. *AXL* is not bound by any MITF family member but is upregulated in the *MITF*/*TFEB* DKO (Figure 6H) but not in the TKO. Upregulation of *AXL* in the *MITF*/*TFEB* DKO is therefore TFE3-dependent.

We also observed many genes in the HALLMARK INTERFERON ALPHA gene set were downregulated in the *MITF* KO but were strongly upregulated by TFE3 in the *MITF*/*TFEB* DKO (Supplemental Figure S6G). As many are bound by MITF family members (Supplemental Table S4) they may be directly regulated with changes in MITF family expression modulating the immune microenvironment.

### MITF, TFEB and TFE3 control tumor growth and immune cell infiltration

Subcutaneous transplantation into in C57BL/6 mice revealed that tumor formation by the *MITF/TFE* family single and double KOs was delayed (Figure 7A-C), with the *MITF* KO, and the DKOs being most affected. Following tail vein injection, lung colonization was severely reduced in the *MITF* KO but was less affected in the *TFEB* and *TFE3* KO cells (Figure 7D, E). Because the immune landscape is dramatically different between human MITF^High^ versus MITF^Low^ tumors^102^, we also examined immune infiltration into the subcutaneous parental and KO tumors. As expected, parental B16-F10 cells exhibited low levels of infiltration of all three immune cell types examined, consistent with the poor immune cell infiltration and immunogenicity of WT B16-F10 tumors (Figure 7F, G). Infiltration by CD8^+^ T-cells increased in the *TFE3* KO and *TFE3*/*TFEB* KO tumors but not in any other KO. This suggests that TFE3 suppresses MITF-dependent CD8^+^ T-cell infiltration.

**Figure 7.**
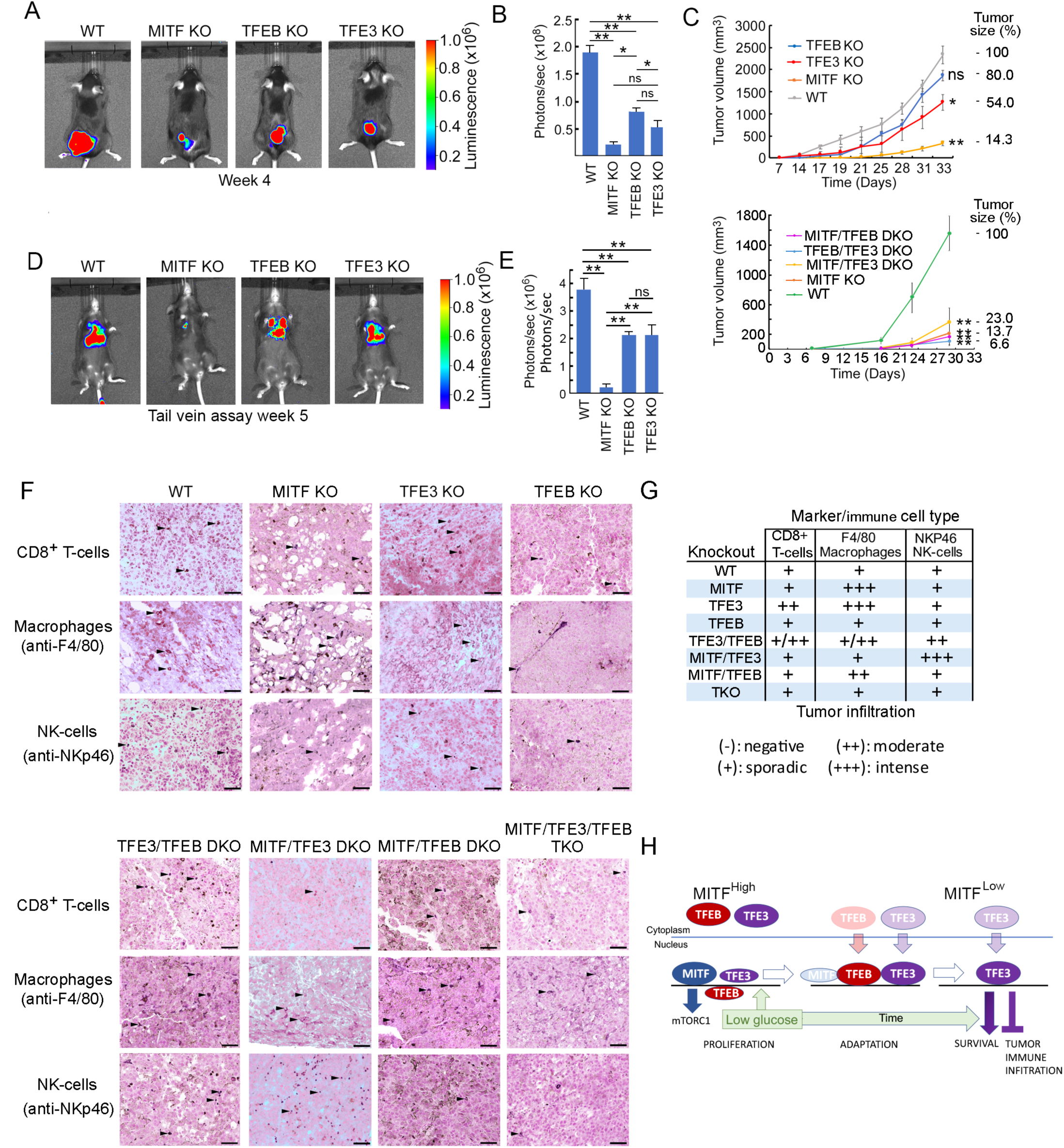
Reduced tumor growth of B16-F10 MITF/TFE KO cells *in vivo*. (A) Bioluminescent imaging of primary tumors 28 days after subcutaneous injection of luciferase-expressing B16-F10 parental or KO cells (n = 5). Firefly luciferin (120 mg/kg) was injected intraperitoneally. Scale bar of bioluminescent image radiance (photons/sec/cm2/sr) color scale was set to Min = 1.00e5; Max = 1.00e6 for all pictures. (B) Quantification of bioluminescence in primary tumors performed using the automatic ROI tool (n= 5) showing average and standard deviation of radiance. *= *p* < 0.05, **= *p* < 0.01 when compared to WT cells using a two-tailed Student t-test. (C) Tumor growth in mice after subcutaneous inoculation of parental or KO B16-F10 cells. n = 5 mice/group. Tumor volume was followed using caliper measurement. *= *p* < 0.05; **= *p* < 0.01; *ns* = not significant when compared to WT cells on the last day. (D) Lung colonization of mice imaged 5 weeks after tail vein injection of 2×10^5^ luciferase-expressing parental or KO B16-F10 cells. N=5 per group. (E) Quantification of bioluminescence in primary tumors performed using the automatic ROI tool (n= 5) showing average and standard deviation of radiance. **= *p* < 0.001, *ns* = not significant using a two-tailed Student t-test. (F) Immune cell infiltration of B16-F10 tumors determined using immunohistochemistry of paraffin embedded sections stained with antibodies to detect T-cells, macrophages and NK-cells. Arrowheads indicate examples of positive cells. Scale bars = 50 μm. (G) Pathologist assessment of immune infiltration in B16-F10 WT and KO tumor sections. Scoring made using 5 fields from 3 tumor sections. (H) Model showing sequential changes in MITF, TFEB and TFE3 expression following glucose limitation over time.

Macrophage infiltration is elevated in tumors derived from the *TFE3* KO, *MITF* KO, and *MITF*/*TFEB* DKO. However, the increase in the *TFE3* KO is moderately reduced if *TFEB* is also knocked out and abrogated in the *MITF/TFE3* DKO. As the TKO shows no increase in macrophage infiltration, the results suggest that TFE3 suppress MITF or TFEB-dependent macrophage infiltration.

No increase in NK-cell tumor infiltration was observed in any of the single KOs or the *MITF*/*TFEB* DKO. However, there is a substantial increase in NK-cells in the *MITF*/*TFE3* DKO tumors, and to some extent the *TFE3*/*TFEB* DKO tumors. As no increase is detected in the TKO, the increase in NK-cell infiltration observed in the *MITF*/*TFE3* or *TFE3*/*TFEB* DKOs is promoted either by *TFEB* or by *MITF* respectively.

Finally, we note that the *MITF* KO tumors, unlike any other, exhibited a different structure with large gaps between clusters of tumor cells that may reflect differential cell-cell adhesion or extracellular matrix composition in the *MITF* KO cells, consistent with a previous report^75^.

## Discussion

Whether functionally related transcription factors with identical DNA binding specificity might perform distinct functions at the same target genes is a key question especially relevant to cancer. Here we use melanoma to examine the role of TFE3 and TFEB in cells that also express the melanocyte-specific isoform of MITF that is largely constitutively nuclear^90^ and not inhibited by mTORC1. The results were surprising.

Consistent with its pro-proliferative role, we found MITF promotes mTORC1 signaling by suppressing expression of *PTEN* and the mTORC1 negative regulators *SESN2* and *DEPTOR*. Signalling downstream from mTORC1 is partitioned into canonical RHEB-dependent signalling to substrates like S6K, and a non-canonical pathway where mTORC1 targeting of TFEB and TFE3 relies instead on RAGC/D and FLCN/FNIP2^9,50,60,92,91,96^. Paradoxically, activating mTORC1 mutations can trigger a negative feedback loop leading to hypophosphorylated nuclear TFEB/TFE3^92,93^. By promoting canonical mTORC1 signaling, it is possible that despite upregulating *RRAGD* and *FNIP2*, MITF may suppress non-canonical signalling to promote the largely nuclear TFEB and TFE3 observed in MITF^High^ melanoma cells. However, surprisingly, TFE3 is cytoplasmic in RAGD-negative IGR39 cells, raising the possibility that rather than cytoplasmic retention, TFE3 might be subject to increased nuclear export.

Remarkably, in response to glucose limitation MITF and then TFEB are downregulated leaving TFE3 to suppress cell death (Figure 7H). The low glucose-mediated switch from TFEB to TFE3 may also be relevant to their activation by AMPK independently of mTORC1^94^. As the switch also operates in colon cancer cells, it seems likely it will occur in many cell types. For example, in macrophages, where TFEB and TFE3 cooperate to control inflammatory signalling^42^, lipopolysaccharide treatment initially activates TFEB before its expression is lost, leaving TFE3 as the sole family member expressed.

Against expectation, while all MITF family members bind the same genes, likely at nucleosome free regions^95,96^, their actions on those genes alone or in combination are not always the same. GSEA suggests that TFEB plays a broadly similar role to MITF, but that TFE3 instead acts in opposition to MITF/TFEB, at least at some genes. Our conclusion is that on encountering microenvironmental stresses, such as glucose limitation, MITF is downregulated via the ISR before TFEB transiently takes on many MITF functions to buy time for cells to restore homeostasis. If the stress remains unmitigated, TFEB is also suppressed, and TFE3 then promotes cell survival, oxidative phosphorylation and upregulation of interferon alpha response genes.

Although MITF is widely assumed to be the key regulator of melanocyte differentiation genes, surprisingly, several differentiation-associated genes were upregulated by TFEB in the *MITF* KO and downregulated if MITF were overexpressed. Notably, in the same *MITF* KO cells, pro-proliferative MITF targets such as *Cdk2* and *Met* were suppressed while MITF-bound anti-proliferative genes such as *Sesn2* and *Deptor* were upregulated. We hypothesize that in proliferative cells MITF activates pro-proliferation genes but suppresses TFEB-driven activation of differentiation genes; by contrast, in differentiated cells, MITF will upregulate differentiation genes and suppress cell division. The nature of this MITF activity switch is unknown but may relate to regulation of MITF co-factors by proliferation-associated post-translational modifications such as acetylation^64,71^.

Previous work revealed that depletion of TFEB in melanoma cells caused reduced proliferation and lower glucose and glutamine uptake, and consequently reduced glycolysis, glutaminolysis and slower tumor growth in vivo^50^. By contrast, Li et al^97^ reported BRAF inhibition increased TFEB-promoted autophagy, while a constitutively active TFEB S142A mutant slowed tumor growth and an S142E phosphomimetic mutant enhanced tumor growth. Our results are more in line with those of Ariano et al^50^ since tumor growth in the B16-F10 *TFEB* KO was moderately reduced, with a greater reduction in any DKO. Our GSEA also revealed an MITF-dependent increase in glycolysis-associated mRNAs in *TFEB* KO cells, perhaps compensating for the decreased glucose uptake and glycolytic enzyme activity observed by Ariano et al^50^ in TFEB-depleted cells. Notably the increased oxidative phosphorylation-dependent ATP production in the *MITF* KO requires TFEB and TFE3 and is consistent with MITF^Low^ slow-cycling melanoma cells exhibiting elevated oxidative phosphorylation^98,99^. Taken together, our data implicate TFEB and TFE3 as playing a key role in the metabolic switch associated with loss of MITF.

Finally, we show that the MITF family controls the quality and quantity of immune infiltration. For example, inactivation of *TFE3* greatly increased MITF-dependent CD8^+^ T-cell and macrophage infiltration, while infiltration by NK-cells was dramatically enhanced in the *MITF*/*TFE3* and *TFE3*/*TFEB* DKOs. The results are consistent with evidence highlighting MITF, and by implication TFE3 and or TFEB, in control of the anti-tumor immune response^80,100,101^. Given the importance of immune infiltration in the response to immune checkpoint blockade, our results provide a clear path to future studies directed at dissecting the molecular mechanisms controlling the tumor immune microenvironment.

### Limitations of this study

The MITF transcription factor family can homo- and heterodimerize^102^ in vitro. However, we cannot establish whether heterodimerization contributes to their capacity to dictate distinct gene expression programs. However, Verastegui et al^103^ found no detectable heterodimerization between MITF and TFE3 in B16-F10 melanoma and genetic analysis of Mitf-family mutant mice indicated heterodimeric interactions are not essential for Mitf-Tfe function^104^. In general, the data are more consistent with expression rather than dimerization playing a key role in their differential activities.

MITF family members bind chromatin-free regions of the genome^95^ with accessibility also dictated by TFAP2^96^. Our ChIP-seq was performed only in WT cells. However, loss of MITF and TFEB could lead to acquisition of novel recognition sites for TFE3, as recently noted^74^. It is possible therefore that some TFE3/TFEB-regulated genes are bound only in the absence of MITF leading to underestimation of the numbers of genes directly regulated by MITF family members.

Finally, KO cell lines are selected for proliferation and consequently cells must bypass any block to proliferation induced by a KO. The assays used here may therefore underestimate effects on the cell cycle by the various MITF-family KOs.

## Resource availability

### Lead contact

Colin R Goding, Ludwig Institute for Cancer Research, Nuffield Department of Clinical Medicine, Old Road Campus Research Building, University of Oxford, Old Road Campus, Headington, Oxford, OX3 7DQ, UK.

### Materials availability

All unique reagents generated in this study may be obtained from the lead contact with a completed materials transfer agreement if necessary.

### Data availability

- All ChIP-seq and RNA-seq datasets have been deposited at the Gene Expression Omnibus (GEO) and are publicly available as of the date of publication. Accession numbers are listed in the Key Resource table.
- Original western blot images have been deposited at Mendeley at doi: 10.17632/p6pr9tv6kv.1 and are publicly available as of the date of publication.
- This paper does not report original code
- Any additional information required to reanalyze the data reported in this paper is available from the lead contact upon request.

## Acknowledgements

We thank Pati Faria-Shayler for technical support. This work was funded by: Ludwig Institute for Cancer Research (CRG, PL, SA, LM); National Institutes of Health R01 CA268597-01 (CRG, PL) and PO1 CA128814-06A1 (YV-G); GABBA, University of Porto (DD); Fundacao de Amparo a Pesquisa do Estado de Sao Paulo (FAPESP) 2017/26148-5 (EO) and 2017/04926-6 (SSM-E); La Caixa (LM); China Scholarship Council (LL); American Cancer Society, via the Holden Comprehensive Cancer Center (HCCC) (IRG-18-164-43); Melanoma Research Alliance (ID#1426690) (CK); Fundación Robles Chillida (Caravaca, Murcia) (L.S-d-C), Ministerio de Ciencia, Innovación y Universidades (MICIU) (Project PID2023-149281OB-100 & FEDER, UE) and Fundación Séneca - Agencia de Ciencia y Tecnología de la Región de Murcia (FSRM/10.13039/100007801 (22544/PI/24) Spain) (JNR-L); Agencia Estatal de Investigación: MCIN/AEI/10.13039/501100011033 PID2023-151128OB-100 and PID2019-10487RB-100 (CG-J) and PID2021-127645OA-100 (ACC); Comunidad de Madrid: PEJ-2021-AI/BMD-22973, PEJ-2021-TL/BMD-21515 and PRECICOLON-CM, P2022/BMD-7212 (CG-J). Fundación Séneca, Región de Murcia (Spain) (21407/FPI/20) (RMD).

## Author contributions

C.R.G. and P.L conceived the project and designed and interpreted experiments. D.D., E.O., S.A, L.M-L, L.L. R.M.D., Y.V-G. and P.L. performed experiments, A.C-C., J.,M.-U. and J.M.G.-M undertook IHC analysis, J.C., P.L. and C.K. provided bioinformatic analysis, P.L., LS-d-C, C.G-J., J.N.R-L. and C.R.G. provided resources and supervision. C.R.G., P.L. C. G-J. and LS-d-C wrote the manuscript.

## KEY RESOURCE TABLE

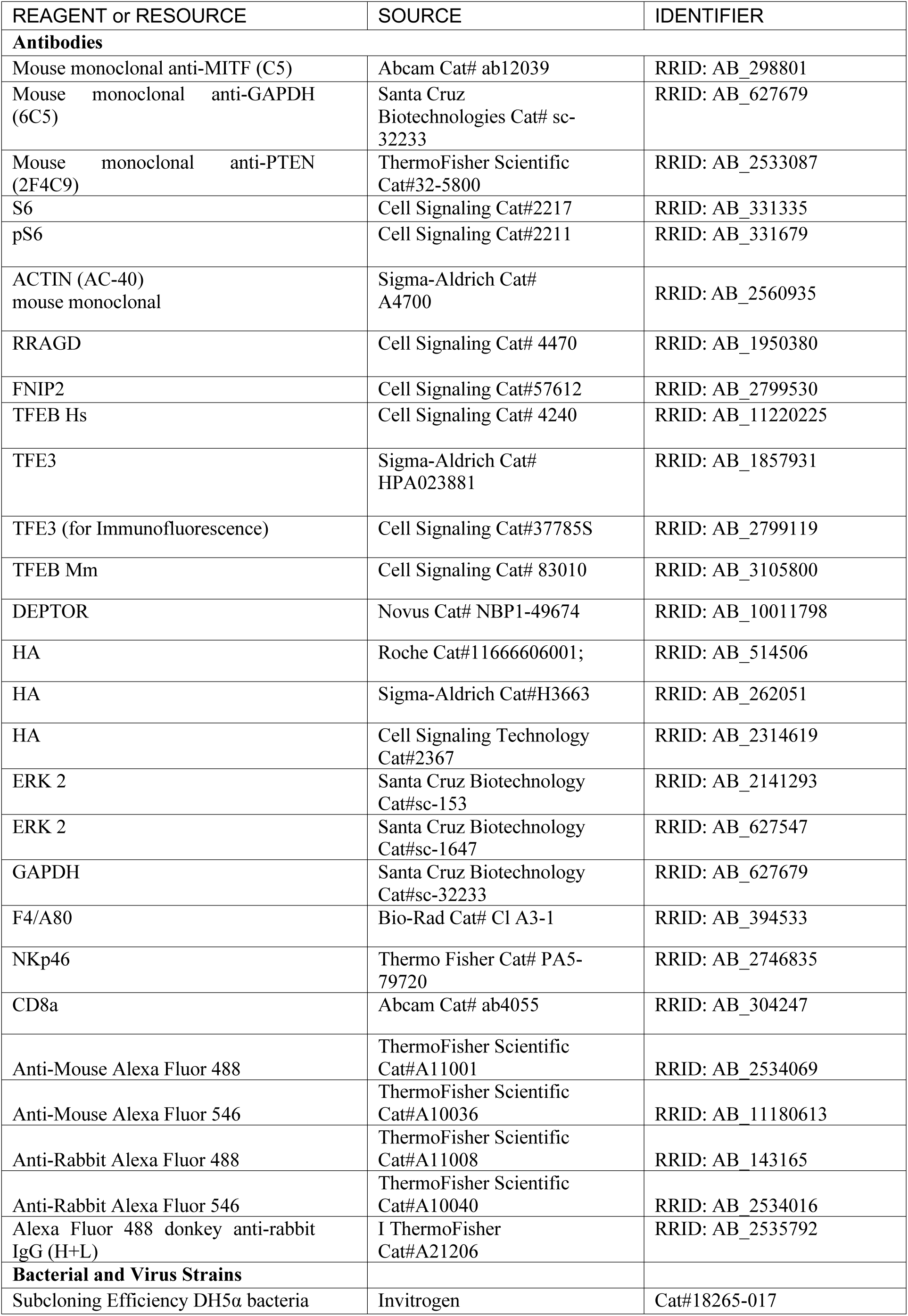
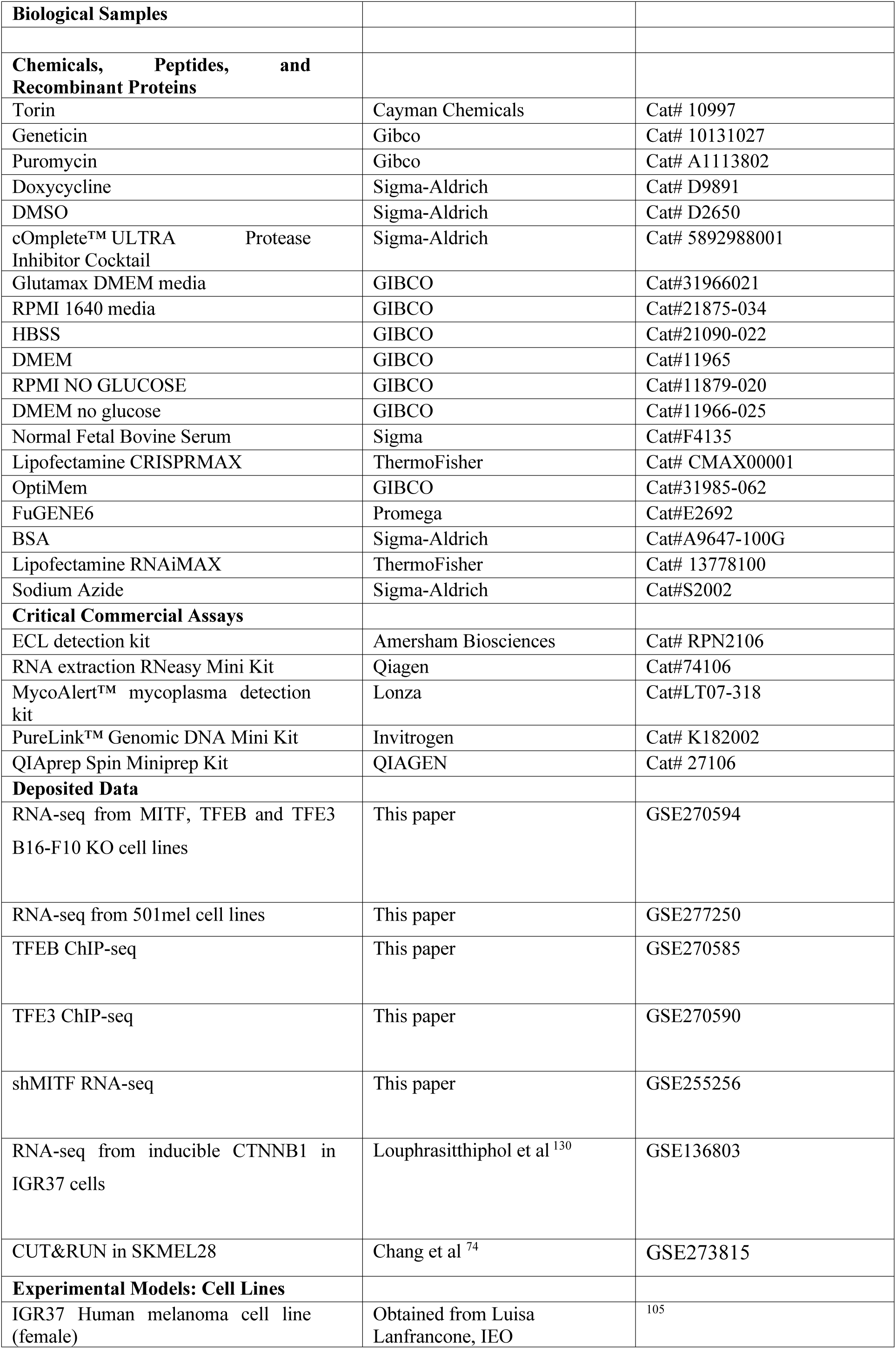
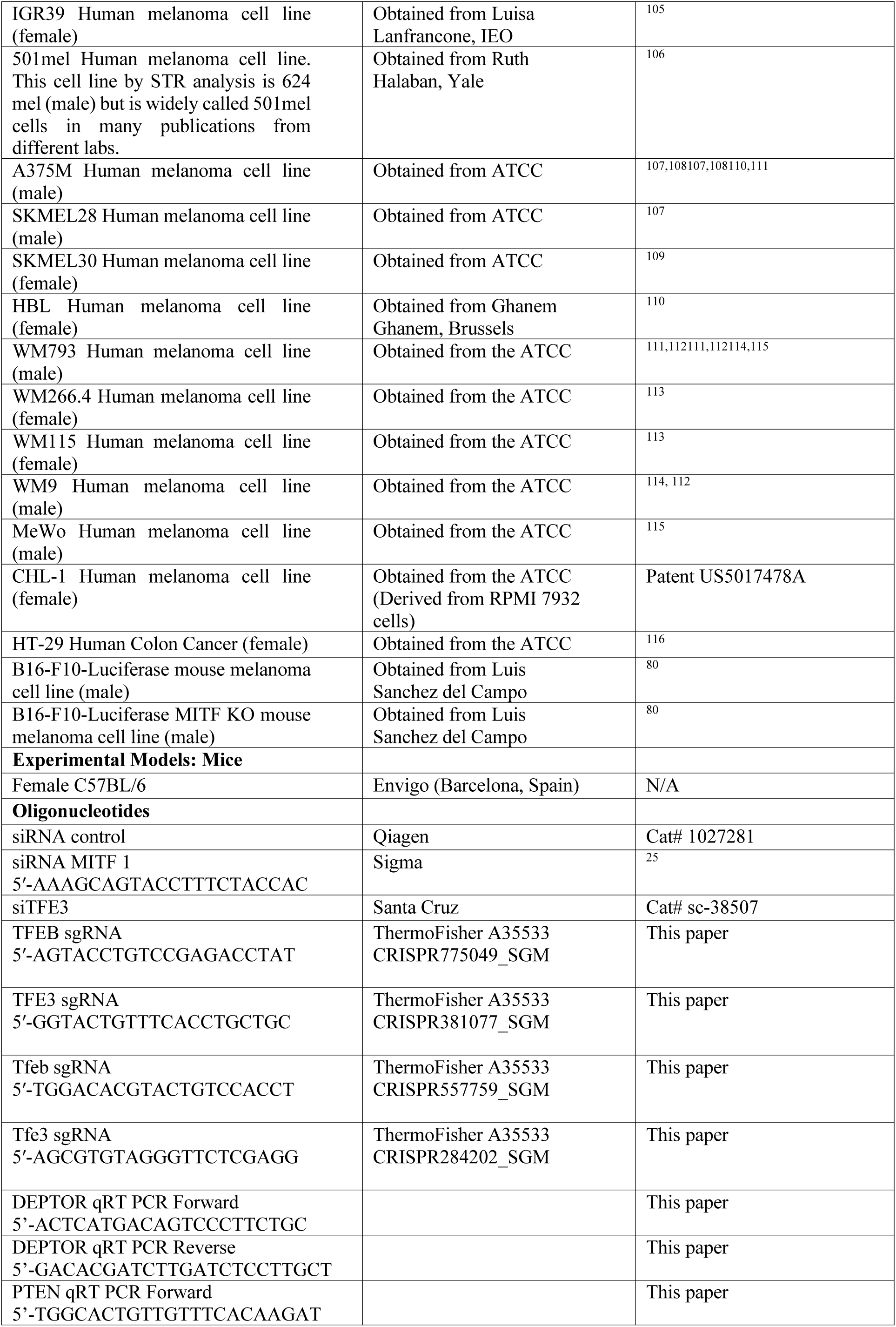
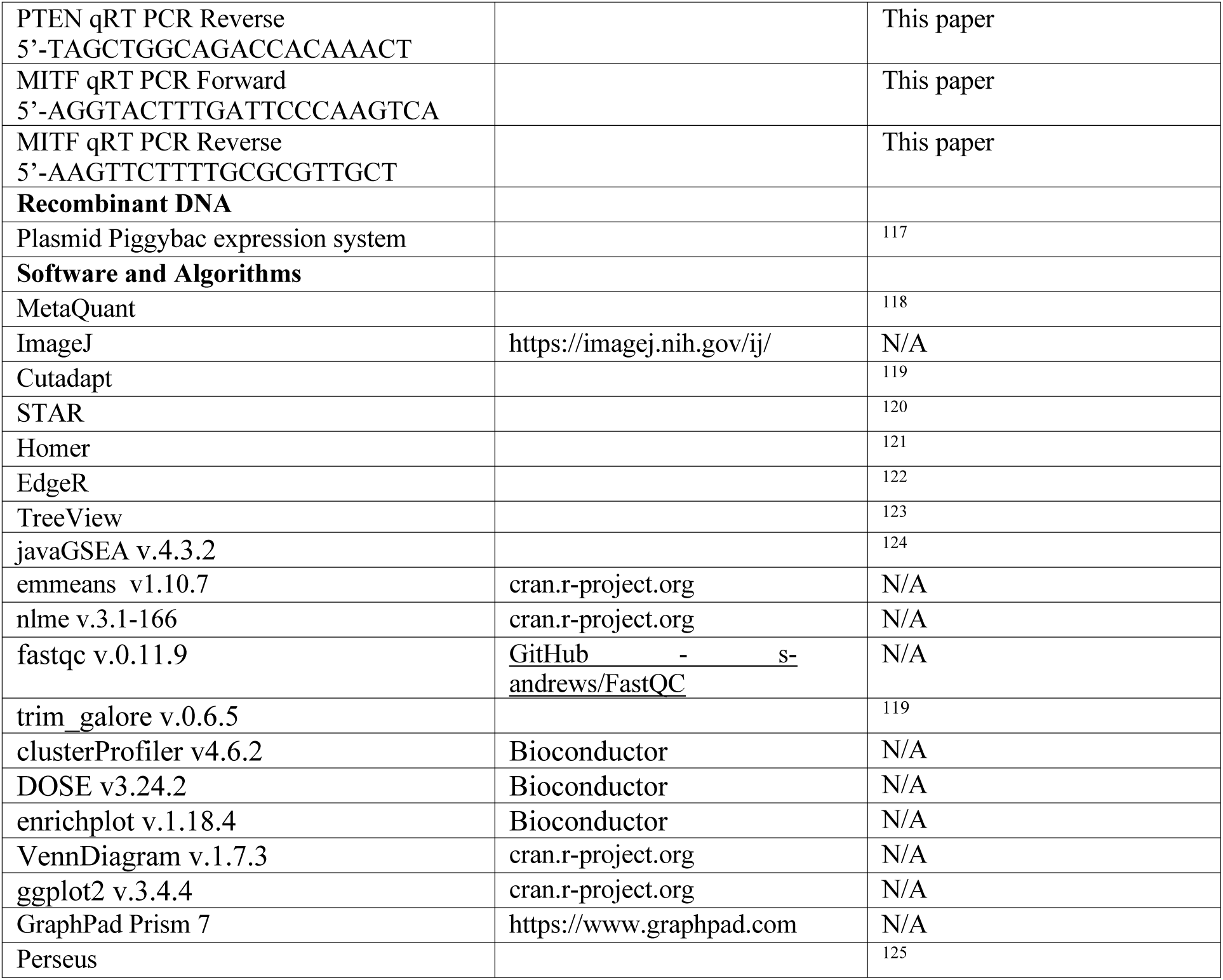

## STAR Methods

### EXPERIMENTAL MODEL AND STUDY PARTICIPANTS

#### Human tumor samples and immunohistochemistry

Melanoma patient samples were obtained from surgical resections of patients diagnosed with melanoma at Hospital Clínico San Carlos (Madrid, Spain). This study using human melanoma tissue was reviewed and approved by the Institutional Review Board (IRB) of Clínico San Carlos Hospital on June 25, 2021 (C.I. 21/498-E). All patients gave written informed consent for the use of their biological samples for research purposes. Fundamental ethical principles and rights promoted by Spain (LOPD 15/1999) and the European Union (2000/C364/01) were followed. Patient data were processed according to the Declaration of Helsinki (last revision 2013) and Spanish National Biomedical Research Law (14/2007, July 3). Patient characteristics are described in the table below. Ethnicity was not recorded.

| Patient Characteristics | N (%) |
| --- | --- |
| **Gender** |  |
| Male | 1 (33.3%) |
| Female | 2 (66.6%) |
| **Age, n(%)** |  |
| < 60 | 1 (33.3%) |
| > 60 | 2 (66.6%) |
| **Breslow (median, mm)** | 2.7 |
| **Clark level, n(%)** |  |
| I | 0 (0%) |
| II | 0 (0%) |
| III | 0 (0%) |
| IV | 3 (100%) |
| **Histologic type, n (%)** |  |
| NM | 2 (66.6%) |
| NA | 1 (33.3%) |
| **Ulceration, n (%)** |  |
| Yes | 1 (33.3%) |
| No | 0 (0%) |
| NA | 2 (66.6%) |
| **Metastasis development** |  |
| Yes | 2 (66.6%) |
| No | 1 (33.3%) |

Paraffin-embedded tumor samples were sliced into 5 μm sections for staining. Slides were deparaffinized by incubation at 60°C for 10 min and then incubated with PT-Link (Dako, Agilent) for 20 min at 95°C at pH 6.0 to detect TFEB. Slides in slide-holders were incubated with peroxidase blocking reagent (Dako, Agilent) and then with the following dilutions of antibodies: anti-TFEB (1:50) overnight at 4°C. Slides were incubated for 20 min with the appropriate HRP-conjugated anti-Ig (EnVision, Dako, Agilent). Sections were then visualized with the HRP Magenta (Dako, Agilent), and counterstained for 5 min with Harrys’ Haematoxylin (Sigma Aldrich, Merck). Photographs were taken with a Zeis microscope with a 20x objective.

#### Mice and mouse tumor immunohistochemistry

Healthy, female C57BL/6 mice, 6 weeks of age, were purchased from Envigo (Barcelona, Spain). All mice were housed in pathogen-free, ventilated cages in the animal facility at the University of Murcia in accordance with the Spanish legislation and directives of the European Community. All animal procedures were approved by the Ethics Committee of the University of Murcia and the Direccion General de Ganaderia y Pesca, Comunidad Autonoma de Murcia (Project reference A13151101). Mice were monitored daily and sacrificed before any signs of distress. Mice were inoculated with 2 × 10^5^ B16-F10 cells subcutaneously or in the tail vein. The size of the subcutaneous tumors was measured three times weekly for the duration of the experiment with calipers. All tumors were analyzed using the IVIS Imaging System (Caliper Life Sciences, Hopkinton, MA, USA). Details of numbers of control and test mice as well as statistical analysis used are provided in the legend to Figure 7.

To establish the degree of inflammatory infiltrate (T-CD8^+^ lymphocytes, NK cells and macrophages), an indirect biotin free HRP immunohistochemistry was carried out in formalin-fixed and paraffin embedded sections from tumors. Briefly, after deparaffination and rehydration, sections underwent a demasking antigen procedure (Dako High pH antigen retrieval solution, Agilent Tech., Barcelona, Spain). Endogenous peroxidase was then blocked using a commercial solution (Dako Peroxidase Blocking, Agilent). Sections were incubated overnight at 4°C with the primary antibodies (polyclonal rabbit anti-CD8, Abcam, Oxford, UK; polyclonal rat anti-F4/80, BioRad, Madrid, Spain and polyclonal rabbit anti-NKp46 (CD335, PA5-79720; ThermoFisher Scientific), and with the anti-rabbit secondary HRP-labelled polymer (Vector ImmPress, Vector Labs., Newark, CA, USA). To avoid false positive staining due to the endogenous melanin pigment located within tumor cells, the immunoreaction was revealed using a special HRP substrate (Vector VIP, Vector Labs.) which labels positive immunoreactions with a dark-violet stain. Finally, sections were contrasted using fast red solution, dehydrated, cleared and mounted. All sections were examined with a standard brightfield microscope (Zeiss Axio A10, Carl Zeiss, Jena, Germany) and representative images were obtained with a high-resolution digital camera (Axiocam 506 color, Carl Zeiss) by using a commercial software (Zeiss Zen Lite, Carl Zeiss). To determine the degree of inflammatory infiltrate a semiquantitative scale was used as follows: - (negative), + (sporadic), ++ (moderate) and +++ (intense) infiltrate.

#### Cell lines

The origin and sex of all cell lines used is provided in the Key Resource table. All cells were subject to authentication using STR analysis using Eurofins authentication service, and routinely tested by mycoplasma via PCR, following a previously established and published protocol ^126^ or use of the MycoAlert PLUS Mycoplasma Detection Kit (Lonza). Only mycoplasma-negative cells were used.

All human melanoma cell lines were grown in Roswell Park Memorial Institute (RPMI) medium, except HBL which were grown in Dulbecco’s Modified Eagle Medium (DMEM), supplemented with 10% Fetal Bovine Serum (FBS), 50 units/mL penicillin and 50 units/mL of streptomycin. All human melanoma cell lines contain the BRAF^V600E^ oncogenic mutation, except HBL and CHL-1 cells which possess wild-type BRAF. The mouse melanoma cell line B16-F10 was grown in DMEM, supplemented with 10% FBS as well as 50 units/mL penicillin and 50 units/mL of streptomycin. All cells were grown as a monolayer and passaged once they reached around 80%-90% confluency.

For cell lines stably transfected with PiggyBac doxycycline-inducible expression vectors, selection and maintenance was performed via the addition of 1 mg/mL of geneticin (G418) and 1 µg/mL of puromycin. Selection was first performed with geneticin for the first week, any live cells then allowed to recover until the second weekly selection treatment with puromycin. Cells were kept at 37 °C with 10% CO2 and grown in T75 culture flasks.

For the *TFEB* and *TFE3* genes the top 10 most efficient sgRNAs in terms of on- and off-site efficacy identified using the Broad Institute Benchling software were evaluated using the Integrated DNA Technologies (IDT) Cas9 Checker software to choose the most efficient gRNA from the list. To generate target gene knockout cell lines, the Lipofectamine CRISPRMAX method (ThermoFisher) was followed. Cells were plated on Day 0 and allowed to adhere in a 6-well plate to reach 35% confluency when transfected on Day 1. 125 µl of Opti-MEM, 6.25 µg of Cas9 nuclease, 1.2 µg of synthetic gRNA and 12.5 µl of Cas9 Plus reagent were added to Tube 1 in that order and allowed to incubate at room temperature. The sequences of the guides were: TFEB 5’-AGTACCTGTCCGAGACCTAT; TFE3 5’-GGTACTGTTTCACCTGCTGC; Tfeb 5’–TGGACACGTACTGTCCACCT; Tfe3 5’–AGCGTGTAGGGTTCTCGAGG. Meanwhile, in Tube 2, 125 µl of Opti-MEM was gently mixed with 7.5 µl of CRISPRMAX reagent and incubated for no more than 3 min. The Lipofectamine mix was then added to the Cas9-gRNA ribonucleoprotein (RNP) complex, mixed well by pipetting, and incubated for 15 min at room temperature. Finally, fresh medium was added to the 6-well plate and the 250 µl lipofectamine-RNP complex mix was added to each well in a dropwise manner. Cells were then incubated a further 24-48 h before screening for monoclonal knockouts. Cells were trypsinized, collected into fresh medium in 5 mL total of suspension, counted using an automated counter and diluted accordingly until reaching about 5×10^5^/ml. At this point, cells were serially diluted into a final 1 ml suspension containing about 1000 cells. Clones were distributed into a 96-well plate, with an estimated density of 0.96 clones per well in a final volume of 200 µl and left to grow for about 2 weeks or until visible colonies arising from single clones were seen. At this stage, cells were trypsinized and expanded until enough material could be obtained for Western Blotting to validate the monoclones. Secondary confirmation of knockout status was performed by extracting genomic DNA from monoclones and sending the PCR-amplified target sequence for Sanger sequencing, provided as a service from Eurofins, to ascertain the genomic event that occurred as a result of the Cas9 nuclease cutting the genome at the gRNA target site.

#### Bacterial strains

For general cloning purposes, the *E.* coli sub-cloning efficiency DH5α competent cells (ThermoFisher Scientific, Cat No. 18265017), were used. The genotype of the DH5α *E. coli* strain is F^-^ Φ80lacZΔM15 Δ(lacZYA-argF) U169 recA1 endA1 hsdR17(r_k_ ^-^, m_k_ ^+^) phoA supE44 thi-1 gyrA96 relA1 λ^-^.

### METHOD DETAILS

#### Cell fractionation

Cells were grown to approximately 70-80% confluency. Subcellular fractionation was performed using the Rapid, Efficient And Practical (REAP) method. Briefly, cells were washed in ice-cold 1X PBS before being scraped into 200 µL of ice-cold 1X PBS containing 0.1% NP-40 (1 tablet of the cOmplete Protease Inhibitor Cocktail (Roche, Cat No. 11697498001) and 1 tablet of the PhosSTOP Phosphatase Inhibitor Cocktail (Roche, Cat No. 4906837001) were added immediately before use) and triturated 5 times using a Gibson P200 pipette. After, 60 µL was removed and transferred to a 1.5 mL Eppendorf tube labelled ‘whole cell fraction’. The remaining 140 µL was centrifuged using a table-top centrifuge (10 sec, 5000xg, 4 °C). After centrifugation, 60 µL of the supernatant was removed and transferred to a 1.5 mL Eppendorf tube labelled ‘cytoplasmic fraction’. The remaining supernatant was removed, leaving only the insoluble pellet. The pellet was washed by resuspending in 200 µL of ice-cold 1X PBS containing 0.1% NP-40 before centrifuging using a table-top centrifuge (10 sec, 5000xg, 4 °C). The supernatant was removed and discarded before resuspending the pellet in ice-cold 1X PBS containing 0.1% NP-40 and labelled ‘nuclear’ fraction. The ‘whole cell’ and ‘nuclear’ fractions were sonicated for 30 sec at 4 °C. After, 20 µL of 4X Laemmli buffer was added to all samples before boiling at 95 °C for 5 min. All samples were stored at -20 °C.

#### Cell cycle and DNA fragmentation analyses

Cells were transfected with 20 nM siRNA (siTFE3 #sc-38507, or siCTRL (QIAGEN # 1027281)), in case of further treatments (-Glc) for another 6 h before samples collected in 1.2 ml DPBS (no Mg^2+^/Ca^2+^) + 5% FBS and fixed in 70% EtOH with gentle shaking at 4°C, 1 h. Cells were stained in 250 µl staining solution (DPBS + 0.5 mg/ml PI, 100 µg/ml (ThermoFisher P3566) RNase A (Roche10109169001), 0.05% Triton X-100) at 37°C, 45’. Following one wash in 1 ml PBS, cells were resuspended in 250-500 µl DPBS + 5% FBS and analysed on flow cytometry (SONY MA900). The cell cycle was profiled using FlowJo (v10.8.1) using Watson model. DNA fragmentation (sub-G1 population) was defined as population with PI signals below the G1 for siTFE3 or directly from Dean-Jett-Fox model fitting.

#### Western blotting

Whole cell lysate was prepared by adding an appropriate amount of 1x Laemmli Sample Buffer (62.5 mM Tris-HCl, pH 6.8, 2% Sodium Dodecyl Sulfate, 10% glycerol, 5% ß-mercaptoethanol, 0.01% bromophenol blue) directly to the cells in the plate, usually 200 µl for one well of a 6-well plate at 80%-90% confluency. The whole lysate was then transferred to a 1.5 mL Eppendorf tube and boiled at 95 °C for 10 min before analysis. 5%-10% of the total sample was used in each lane.

Proteins were analyzed via sodium dodecylsulphate polyacrylamide gel electrophoresis (SDS-PAGE) using 10% acrylamide from a 30% stock solution with 37.5:1 acrylamide to bis-acrylamide ratio (Severn Biotech). To allow fine resolving of specific phosphorylated proteins from their non-phosphorylated counterpart, an acrylamide stock with a ratio of 200:1 acrylamide to bis-acrylamide was used (Severn Biotech).

Samples were run/separated at 140V for 90 min in 1x Running Buffer (25 mM Tris, 192 mM glycine, 0.1 % SDS). Precision Plus Dual-Colour Standard (BioRad) was used as indicated by the manufacturer was used as a size marker. After SDS-PAGE proteins were transferred unto a nitrocellulose membrane (GE Amersham) in a wet transfer tank (BioRad) filled with 1x Transfer Buffer (25 mM Tris, 192 mM glycine, 20 % ethanol) and transferred at a constant amperage of 400 mA for 75 min. Membranes were then placed in 50 mL Falcon tubes and blocked at room temperature for 60 min in 5% Bovine serum albumin (BSA) in Tris buffered saline (TBS) containing 0.1% Tween (TBS-T). Membranes were then briefly rinsed with TBS-T before proceeding to primary antibody incubation, also performed in 5% BSA in TBS-T. Most primary antibodies were used in a 1:1000 dilution while housekeeping antibodies were diluted 1:5000. Overnight incubation with primary antibodies was performed by leaving the 50 mL Falcon tubes rolling and incubating in the cold room at 4 °C. The following day, membranes were rinsed twice for 10 min before incubation with HRP-conjugated secondary antibodies diluted 1:5000 in 5% milk in TBS-T at room temperature for 120 min. Afterwards, membranes were again rinsed and washed 2 times for 10 min, at which point enhanced chemiluminescence reagent (ECL, GE Amersham) was incubated on the membranes for 5 min before the membranes were exposed to X-ray film (Fujifilm) and developed.

#### Immunofluorescence

For immunofluorescent experiments, cells were plated on circular glass coverslips (VWR) and grown until reaching 70-80% confluency in 24-well plates. Medium was aspirated and cells washed with 1x PBS at room temperature and then fixed on the plate with 4% paraformaldehyde (PFA) in 1x PBS for 10 min at room temperature. After fixation, PFA was aspirated, and coverslips rinsed 3 times with PBS followed by permeabilization with 0.2% Triton X-100 in PBS at room temperature for 10 min with agitation on a shaker. Permeabilization buffer was then aspirated, and fresh PBS was added to the wells to rinse the fixed cells on the coverslips before blocking for 1 h at room temperature with 4% BSA in PBS in a shaker. Afterwards, coverslips were, dried and placed face down on a PARAFILM M (Sigma) sheet with 50 µl of primary antibody solution, consisting of primary antibody normally diluted 1:100 in 4% BSA in 1x PBS. Antibody incubation was either performed for 2 h at room temperature or overnight at 4° C, depending on affinity, concentration and ultimate signal obtained using the primary antibody. Coverslips were then placed into in a new sterile 12-well plate and washed 3x with PBS for 5 min each, before simultaneous incubation with secondary antibodies and DAPI, diluted both 1:500, face down in the parafilm with a light-protective cover. Once the secondary incubation was complete, coverslips were again transferred to a plate and washed with PBS for 5 min 3x times, dried and finally mounted on glass microscope slides (VWR) with MOWIOL (Millipore) mounting medium. Images were then taken with a confocal Zeiss LSM710 or LSM980, with image acquisition performed with the matching proprietary Zen software (Zeiss). Image processing was for presentation and publication purposes was done via Photoshop. Antibodies used are listed in Key Resource table.

#### Proliferation/survival assays

A DNA content-based cell proliferation assay that is independent of cellular metabolic status was carried out using CyQUANT™ Direct Cell Proliferation Assay (ThermoFisher, # C35011) as per manufacturer protocol using media only for background subtraction.

#### Seahorse analysis

The oxygen consumption rate (OCR) and extracellular acidification rate (ECAR) were measured using a Seahorse XF96 extracellular flux analyzer (Seahorse Bioscience, North Billerica, MA, USA). Briefly, 7,000-14,000 B16-F10 cells were seeded in XF96 well plates with the objective of achieving approximately 75% confluence and homogeneous distribution. On the following day, the medium was replaced with fresh Seahorse XF medium without phenol red supplemented with 1 mM pyruvate, 2 mM glutamine, and 10 mM glucose and incubated one hour in a CO2-free incubator to guarantee precise measurements of extracellular pH. Metabolic activity was analyzed using an Agilent Seahorse 96XF device and respective kits. Glucose pyruvate oxidation stress, Glutamine oxidation stress and Glycolytic rate tests were performed as described in the manufacturer’s protocols respectively (Kit 103673–100, Kit 103674–100 and Kit 103344–100). Respiratory parameters were calculated using the Seahorse Wave Desktop Software (Agilent Technologies).

#### RNA-seq

Cells were allowed to reach 80% confluency in a 6-well plate before total RNA was extracted using RNeasy kit (QIAGEN #74106) as per manufacturer protocol. 500 ng total RNA were used for library preparation using QuantSeq Forward kit (LEXOGEN #015.96) according to the manufacturer’s protocol before sequencing by the Wellcome Trust genomic service, Oxford.

#### ChIP-seq and CUT&RUN

The ChIP protocol used here was the same as that previously used for HA-MITF ChIP seq. Cells were grown in 15 cm^2^ dishes to reach 70-80% confluency with each replicate making up to 30 dishes. Cells were washed with 1x PBS and trypsinised with 4 mL of trypsin before being collected with 20 mL of warm medium into a 50 mL Falcon placed on ice. Cells were then centrifuged at 800 g for 4 min at 4 °C. The medium was aspirated, and the cell pellet resuspended in 1 mL of ice-cold 1x PBS, to which 35 mL of ice-cold 1x PBS 0.4% PFA was added for crosslinking proteins to the DNA. The Falcon tubes were then kept on a roller for 10 min at room temperature before the crosslinking reaction was stopped by adding 4 mL of 2 M Glycine (0.2 M final concentration) to quench the PFA for a further 10 min at room temperature on the roller. Tubes were then centrifuged at 1500 g for 10 min at 4 °C and the supernatant removed. At this stage, the cell pellets were frozen at -80 °C. Then 1 ml of fresh ChIP lysis buffer (50 mM Tris-HCl (pH 8.0), 10 mM EDTA, 10 mM sodium butyrate, 1% SDS, 4×PIC (Roche; Cat#05056489001)) was added to the pellets. PIC and PhosStop were added to the buffer just immediately prior to use. Replicate lysates were pooled and broken down by using a 25 G needle and a 20 ml syringe, and aliquoted into Covaris 1 ml vials that were sonicated in a S220 sonicator (Covaris) for 15 min at 160 W, with 5% duty cycle and 200 cycles per burst. Sonicated DNA was then transferred from the Covaris vessel to a 1.5 ml Eppendorf tube and centrifuged at 13,000 g for 10 min at 4 °C. The cleared supernatant was then transferred to a fresh 1.5 ml Eppendorf tube stored at -80°C after two 10 µl aliquots were transferred to a new tube. One aliquot was kept as a 1% input control and frozen at -80 °C. The second aliquot was reverse crosslinked overnight to check for DNA fragment size by running the purified sample on a 2% agarose gel to confirm successful sonication and a desired size of fragment size of 200-400 bp. Once correct fragmentation had been achieved, DNA samples were thawed at 4 °C and diluted with ChIP dilution buffer to 9 ml in a 50 ml Falcon tube. Next, 120 µg of HA antibody (Roche) was used per 30 dishes and the 50 ml Falcons left rotating and incubating overnight in the cold room at 4 °C. In parallel, using an Eppendorf magnetic rack, 600 µl of Protein G Dynabeads were washed, resuspended and ultimately blocked overnight in a 1.5 mL tube with ChIP dilution buffer containing 0.5 mg/ml of BSA (Sigma) at 4°C in a rotator in the cold room. For the pulldown, blocked Dynabeads were added to the sample and rotated for 60 min at 4°C in the cold room, before being centrifuged for 10 min at 1500 g and 4 °C. The supernatant was removed and all further washing steps carried out with the use of a microcentrifuge tube magnetic rack in the 4 °C cold room, with the beads transferred to a fresh new tube after each washing step. Beads were resuspended in 1 ml of ChIP low salt wash buffer and transferred to a new 1.5 ml microcentrifuge tube, washed again and resuspended in another 1 ml of ChIP low salt wash buffer before being placed in a new tube. Beads were then washed twice more with ChIP low salt wash buffer followed by 2x washes with ChIP high salt wash buffer, then 2x washes with LiCl wash buffer. Finally, ChIP samples were simultaneously eluted with freshly made microwaved elution buffer and reverse cross-linked overnight in a Eppendorf shaker set at 55 °C and 600 rpm, to release the proteins from the beads and the DNA fragments from the proteins. ChIP DNA was recovered using a PCR purification kit according to the manufacturer’s instructions, with the caveat that the first elution step was made with 30 µl for ChIP-seq library preparation and the second elution with 120 µl was kept for ChIP-qPCR. ChIP and Input samples were tested on Qubit and Bioanalyzer to confirm amount and integrity of the samples before being sent for sequencing CUT&RUN data for TFE3 and MITF in SKMEL28 WT and MITF KO cells was generated by Chang et al^74^ from the MITF KO cell lines established by Dilshat et al ^75^.

#### Bioinformatics analyses

B16-F10 RNA-seq and TFE3 ChIP-seq libraries were sequenced on an Illumina NovaSeq 6000. TFEB ChIP-seq libraries were sequenced on an Illumina HiSeq 4000. 501mel RNA-seq were sequenced on NextSeq 550. Base calling and demultiplexing were carried out by the Oxford Genomic Centre Service, University of Oxford.

Fastq reads for each sample were sequenced across multiple flow cells (except B16-F10 RNA-seq libraries which were sequenced on a single flow cell) and were stitched on UNIX and quality controlled using fastqc (v.0.11.9).

ChIP-seq samples were adaptors and 2 colour-trimmed as paired-end reads while RNA-seq were also trimmed of poly-A as single-end reads using trim_galore (v.0.6.5; running python-base v.3.6.10, cutadapt v.2.10, java v.17.0.1, fastqc v.0.11.9 and pigz v.2.4)^119^.

B16-F10 trimmed reads were mapped to the *Mus musculus* reference genome (mm39) with STAR (v.2.7.8a)^120^. 501mel trimmed reads were mapped to the Human reference genome (hg38) with STAR (version c.2.7.8a). RNA-seq normalisation and differential gene expression analyses were performed as previously described^18,19,127^ using edgeR glmQLFTest. GSEA analyses were performed using pre-ranked lists derived from edgeR glmQLFTest -log10(P-value)/sign(logFC) using GSEA (v.4.3.2) against MSigDB v2022.1. Dot blot of GSEA analyses were generated using pre-ranked list above using the Bioconductor packages clusterProfiler (v4.6.2), DOSE (v3.24.2). Genesets from MSigDB were imported using the package msigdbr (v7.5.1).

TFEB and TFE3 ChIP-seq trimmed fastq files were mapped using STAR (v.2.5.1b or v.2.7.8a, respectively)^120^ against the human hg38 index. Peak calling, *de novo* motif identification, genome ontology analyses were performed using Homer as previously described^71,127^. Normalised bigwig files of aligned reads were generated using Homer for visualisation on the UCSC genome browser as MultiWig hubs. The peaks calling parameter were: FDR rate threshold = 0.001, Fold over input = 4.00, Poisson p-value over input = 1.00e-04, Fold over local region = 4.00, Poisson p-value over local region = 1.00e-04 and variable parameters (modelled according to each dataset as these are sequencing depth dependent), FDR effective Poisson threshold and FDR tag threshold

#### Data visualisation

Overlapping peaks were visualised using a R package VennDiagram (v.1.7.3). Genome ontology, box and jitter plots, volcano plots and peak correlation plots were generated using an R package ggplot2 (v.3.4.4). Heatmaps and hierarchical clustering of normalised gene expression were performed using an R package pheatmap (v.1.0.12). Tag density of ChIP-seq datasets were visualised using the java package TreeView (v.1.1.6) ^123^.

GSEA dot plots were generated using the Bioconductor package enrichplot (v.1.18.4) using the cut-off adjusted-p-value of 0.05 (Benjamini-Hochberg procedure).

Moving average plots were generated using TCGA log2 of RSEM transcript abundance quantification using 20 samples window. Spearman’s *ρ* computation and correlation tests were performed using R cor.test function.

### QUANTIFICATION AND STATISTICAL ANALYSIS

Details of sample numbers, statistical analyses and significance are provided in the Figure legends. Figure 1A: Error bars indicate S.D., n = 3, **= p <0.01.

Figure 1D: The moving average expression trendline for *RRAGD* or *FNIP2* were made using a sample window size of n=20. Spearman correlation coefficient was used to test for association between paired samples, *RRAGD* or *FNIP2* with *MITF*.

Figure 1G-H: n = 3. The box and whisker plots show the median, two hinges corresponding to 25^th^ and 75^th^ percentiles and two whiskers extending 1.5× IQR. The black lines indicate means. For Pairwise comparisons between control (-Dox) and test (+Dox) all p = <0 .0001calculated using edgeR glmQLFTest.

Figure 2A-C: The moving average expression trendline for *TFE3* or *TFEB* were made using a sample window size of n=20. Spearman correlation coefficient was used to test for association between paired samples, *TFE3* or *TFEB* with *MITF* and *TFE3* with *TFEB*.

Figure 2I: Error bars indicate S.D. N=3. *= p <0.05; **= p <0.01; ***= p <0.001; ****= p <0.0001 ns = not significant, Student t-test. No. of cells analysed siCTRL DMEM (206,097, 200,000 and 200,000); siCTRL -Glucose (200,000, 200,000 and 200,000); siTFE3 DMEM (200,000, 200,000 and 125,305); and siTFE3 -Glucose (200,000, 200,000 and 52,884)

Figure 3C-J

All peaks were called using the following fixed parameters

\# FDR rate threshold = 0.001000000

\# Fold over input = 4.00

\# Poisson p-value over input = 1.00e-04

\# Fold over local region = 4.00

\# Poisson *p*-value over local region = 1.00e-04

and variable parameters (modelled according to each dataset as these are sequencing depth dependent)

\# FDR effective poisson threshold

\# FDR tag threshold

Figure 3C: The box and whisker plots show the median, two hinges corresponding to 25^th^ and 75^th^ percentiles and two whiskers extending 1.5× IQR. The black lines indicate means.

Figure 3G: Boxes indicate interquartile range (25^th^ – 75^th^ percentile) and median (50^th^ percentile). The whiskers span minimum and maximum values within 1.5 × IQR. **** indicates p-value < 0.001

One-way ANOVA.

Figure 3H: *De novo* motif identification using HOMER. The statistic accounts for multiple hypothesis testing across all candidate motifs. E-value=P-value×number of tested motifs. Where P-value denotes the probability that the observed enrichment (or greater) occurs by chance; the E-value denote the expected number of motifs with equal or better enrichment by chance.

Figure 3I: Enrichment is calculated using Homer cumulative hypergeometric test, plot as - LN(Enrichment). % overlapped were calculated from peaks overlapping the feature divided by total peaks called.

Figure 4A: Heatmap of normalized gene expression of differentially expressed genes between 501mel parental and HBSS treated (edgeR glmQLFTest, FC ≥ 2, p = <0.05).

Figure 4B: Volcano plot of differential gene expression (edgeR glmQLFTest, FC ≥ 2, p = <0.05).

Figure 4C: GSEA dot plots of pre-ranked values calculated from edgeR glmQLFTest -log10(P-value)/sign(logFC) using the cut-off adjusted-p-value of 0.05 (Benjamini-Hochberg procedure).

Figure 4F: Heatmap of normalized gene expression of differentially expressed genes between indicated pairs (edgeR glmQLFTest, FC ≥ 2, p = <0.05)

Figure 4G: Volcano plots of differential gene expression between indicated pairs (edgeR glmQLFTest, FC ≥ 2, p = <0.05).

Figure 4H: GSEA dot plots of pre-ranked values calculated from edgeR glmQLFTest -log10(P-value)/sign(logFC) of the indicated pair, using the cut-off adjusted-p-value of 0.05 (Benjamini-Hochberg procedure).

Figure 5C: Heatmap of normalized gene expression of differentially expressed genes between indicated samples relative to parental control (edgeR glmQLFTest, FC ≥ 2, p = <0.05).

Figure 5D-K: GSEA dot plots of pre-ranked values calculated from edgeR glmQLFTest -log10(P-value)/sign(logFC) of the indicated pair, using the cut-off adjusted-p-value of 0.05 (Benjamini-Hochberg procedure).

Figure 6A-D: For Seahorse assays, values were normalized to cell number. For the quantification and statistical analysis three or more independent repeats were performed for each experiment unless stated otherwise in the figure legend. The data were analyzed using SPSS statistical software for Microsoft Windows, release 6.0 (Professional Statistic, Chicago, IL, USA). For comparison of two groups two-tailed Student t-test was used. Mann Whitney-U test was used for comparison of two groups with non-normal distribution. Values were considered significantly different when *p*<0.05.

Figure 6A: Error bars = SD; n=3

Figure 6B-C: Error bars =SD; n=3; ns = non-significant, *= p<0.05, **= p<0.01 (black, comparison with parental cells; red, comparison with TKO); Student t-test.

Figure 6D: Numbers indicate % derived from each source. Error bars =SD; N=3; ns = non-significant, *= p<0.05, **= p<0.01, Student t-test comparing to parental WT B16-F10 cells.

Figure 6E-I Heatmap of normalized gene expression (CPM calculated using edgeR) at indicated genes.

Figure 7A-B: n =5, bar show average and standard deviation of radiance. *= p < 0.05, **= p < 0.01 using a two-tailed Student t-test.

Figure 7C: n =5, *= p < 0.05; **= p < 0.01; ns = not significant when compared to WT cells on the last day using a two-tailed Student t-test.

Figure 7D-E: n =5, bar show average and standard deviation of radiance. **= p < 0.001, ns = not significant using a two-tailed Student t-test.

Figure S2B, G: Spearman correlation coefficient was used to test for association and calculate Spearman’s p-value between paired samples for both replicates.

Figure S2E, H, J: Reference peaks were called using the same parameter as Figure 3C-J against indicated input.

Figure S3A: Heatmap of normalized gene expression of differentially expressed genes between indicated samples relative to parental control (edgeR glmQLFTest, FC ≥ 2, p = <0.05).

Figure 3B: Volcano plots of differential gene expression between indicated pairs (edgeR glmQLFTest, FC ≥ 2, p = <0.05).

Figure S3C-F: GSEA dot plots of pre-ranked values calculated from edgeR glmQLFTest -log10(P-value)/sign(logFC) of the indicated pair, using the cut-off adjusted-p-value of 0.05 (Benjamini-Hochberg procedure).

Figure S4A-H: GSEA dot plots of pre-ranked values calculated from edgeR glmQLFTest -log10(P-value)/sign(logFC) of the indicated pair, using the cut-off adjusted-p-value of 0.05 (Benjamini-Hochberg procedure).

Figure S4J: N=12 (3 biological replicates in quadruplicate). Error bars indicate S.D. **= p < 0.01; **** = p < 0.0001; ns = not significant when compared to WT cells at 96 h. Linear Mixed-Effects Models was used first to model the slopes then a post-hoc comparison using estimated marginal means accounting for two interacting factors (cell line and time) was used for pairwise comparison. The Bonferroni method was used to adjust p-values for multiple comparisons.

Figure S5A: Heatmap of normalized gene expression of differentially expressed genes between indicated samples relative to parental control (edgeR glmQLFTest, FC ≥ 2, p = <0.05) filtering out for genes that are commonly bound by all three factors.

Figure S5B: The p-value for each overlap is computed with the hypergeometric distribution (equivalently, one-tailed Fisher’s exact test), and the FDR q-value is obtained by applying the Benjamini–Hochberg (BH) multiple-testing procedure across all gene sets tested.

Figure S6B, G: Heatmap of normalized gene expression (CPM calculated using edgeR) at indicated genes.

Figure S6F: N=3. Error bars = S.D. **** = p < 0.0001.

## Supplemental information

Document S1. Figures S1-S6.

Supplemental Table S1: DEGs in 501mel cells treated with HBSS or Torin, or 501mel KO cells, related to Figure 4 and Supplemental Figure S3.

**A)** + Torin vs Control
**B)** HBSS vs Control
**C)** *TFE3* KO vs Parental
**D)** *TFE3* KO + Torin vs Parental +DMSO
**E)** *TFE3* KO +Torin vs Parental +Torin
**F)** *TFEB* KO vs Parental
**G)** *TFE3* KO + Torin vs Parental +DMSO
**H)** *TFE3* KO +Torin vs Parental +Torin
**I)** *TFE3/TFEB* DKO vs Parental
**J)** *TFE3/TFEB* DKO vs Parental +DMSO
**K)** *TFE3/TFEB* DKO vs Parental +Torin

Supplemental Table S2: DEGs in B16-F10 KO cells, related to Figure 5 and Supplemental Figure S4.

**A)** *MITF* KO vs Parental
**B)** *TFE3* KO vs Parental
**C)** *TFEB* KO vs Parental
**D)** *MITF*/*TFE3* DKO vs Parental
**E)** *MITF*/*TFEB* DKO vs Parental
**F)** *TFE3*/*TFEB* DKO vs Parental
**G)** *MITF*/*TFE3*/*TFEB* TKO vs Parental

Supplemental Table S3: DEGs in B16-F10 KO cells commonly bound by all MITF family members, related to Figure S5A.

**A)** Numbers of genes in each cluster
**B)** Genes in Cluster 1
**C)** Genes in Cluster 2
**D)** Genes in Cluster 3
**E)** Genes in Cluster 4
**F)** Genes in Cluster 5
**G)** Genes in Cluster 6
**H)** Genes in Cluster 7
**I)** Genes in Cluster 8
**J)** Genes in Cluster 9.1
**K)** Genes in Cluster 9.2
**L)** Genes in Cluster 10.1
**M)** Genes in Cluster 10.2

Supplemental Table S4: HALLMARK Interferon Alpha Response genes bound by MITF family members, related to Figure S6.

---

**Figure S1 related to Figure 1.**
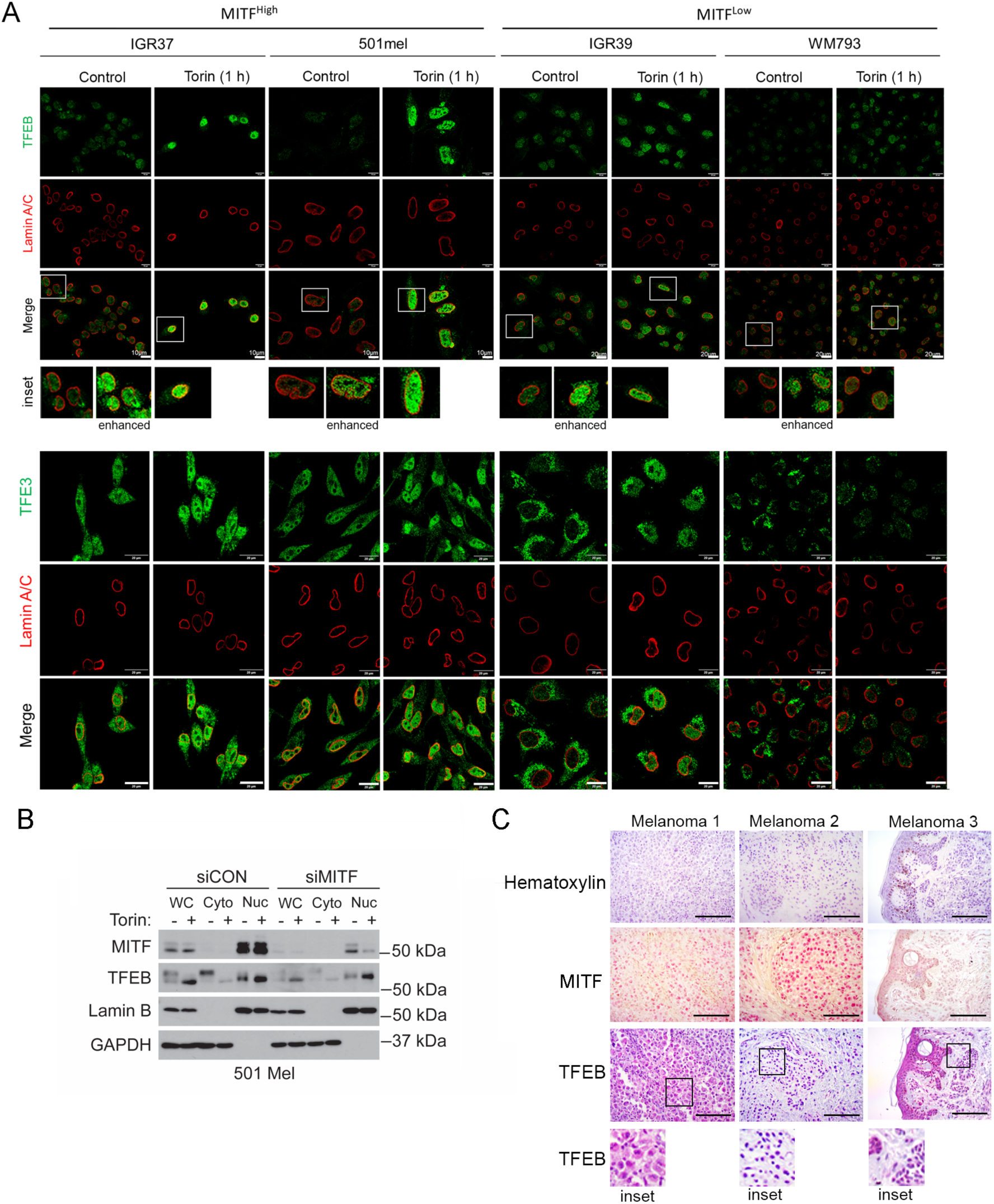
Predominantly nuclear localization of TFEB in melanoma cells. (A) lmmunofluorescence in indicated melanoma cell lines treated or not with 250 nM Torin for 1 h using anti-TFEB (green; top panels), anti-TFE3 (green; bottom panels) and anti-Lamin B (red). Scale bars= 10 µm for 501mel and IGR37, and 20 µm for the IGR39 and WM793 cells for the TFEB panels and 20 µm for the TFE3 panels. Insets for TFEB panels show magnified regions below, and for the control samples an enhanced image is shown to enable to nuclear localization of TFEB to be readily visualized. (**B**) Western blot showing fractionated 501mel cell extracts using Lamin B and GAPDH as markers for the nuclear and cytoplasmic fractions. Cells were transfected with control or siMITF for 48 h before treatment with 250 nM Torin or DMSO before fractionation. (C) lmmunohistochemistry on three human melanomas using anti-TFEB antibody (lower panels) and hematoxylin staining (upper panels). Scale bars= 50 µm.

**Figure S2 related to Figure 3.**
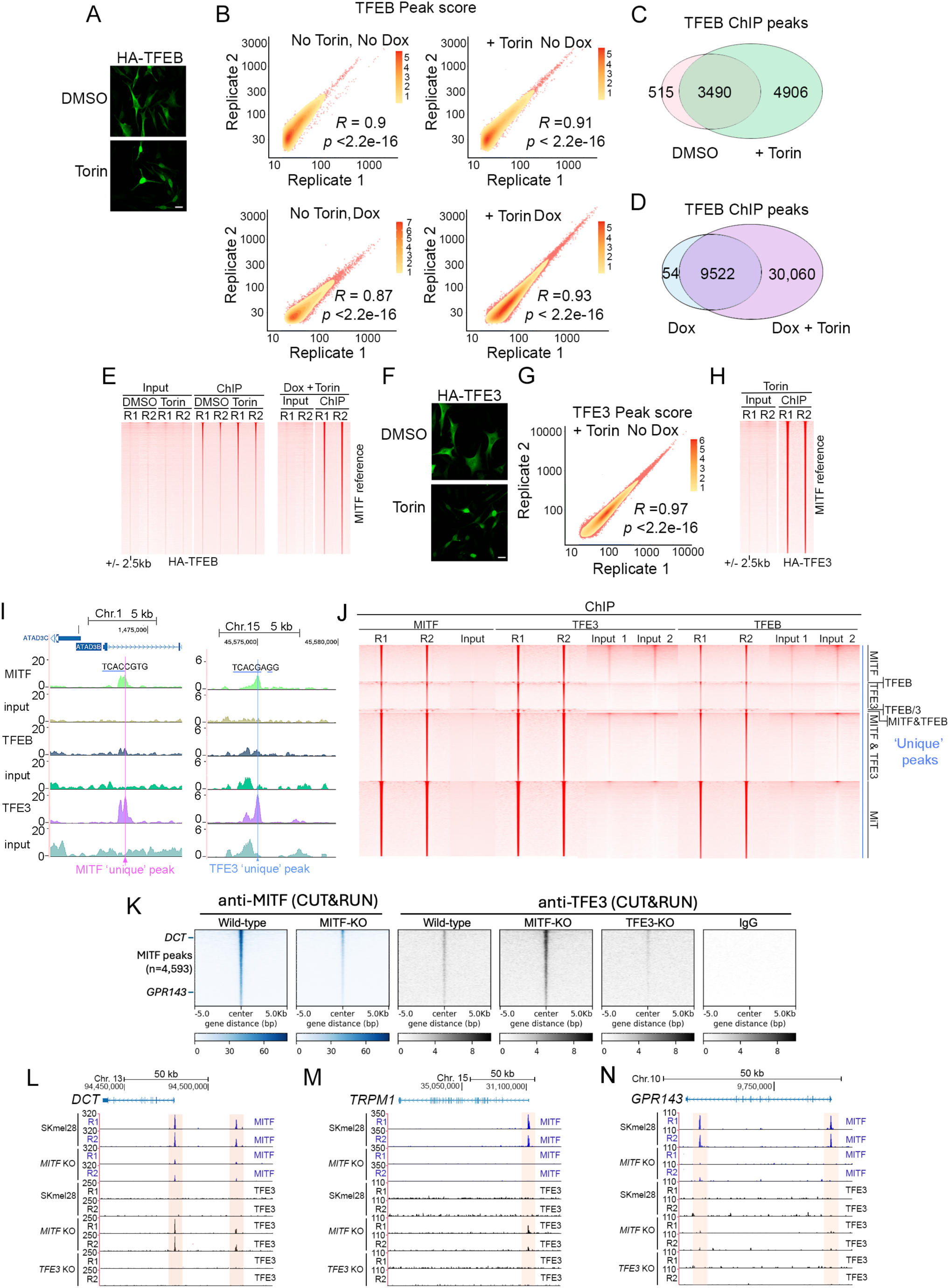
ChlP-seq of HA-TFEB and HA-TFE3. (A) lmmunofluorescence using anti-HA antibody showing nuclear accumulation of HA-TFEB after treatment with 250 nM Torin for 6 h. Scale bar= 10 µm. (B) Inter-replicate correlation of ChlP-seq of HA-TFEB performed under indicated conditions. R = Spearman rho. (C-D) Venn Diagrams showing significant HA-TFEB ChlP-peaks from 501mel cells with or without Torin treatment (C) or in the presence of 5 ng Doxycycline with or without 250 nM Torin (D). (E) Heatmap of tag density of each HA-TFEB ChlP-seq experiment using O or 5 ng doxycycline with or without 250 nM Torin as indicated using ranked MITF ChlP-seq narrow peak as reference genome coordinate and extending +/- 2.5kb. (F) lmmunofluorescence using anti-HA antibody showing nuclear accumulation of HA-TFE3 after treatment with 250 nM Torin for 6 h. Scale bar= 10 µm. (G) Inter-replicate correlation of ChlP-seq of HA-TFE3 performed under indicated conditions. (H) Heatmap showing tag density of each HA-TFE3 ChlP-seq experiment of HA-TFE3 expressed using O ng doxycycline with 250 nM Torin using the same ranked MITF narrow peaks reference as in E. (I) UCSC genome browser screenshots showing ChlP profiles for input control and HA-TFEB, HA-TFE3 and HA­ MITF at indicated loci called as an MITF ‘unique’ peak (left) and TFE3 ‘unique’ peak (right). Only one replicate is shown. Similar results were obtained for replicate 2. Sequences related to the MITF family binding consensus are shown above the tracks with the consensus-related sequence underlined. (J) Heatmap of tag density of each HA-MITF, HA-TFEB and HA-TFE3 ChlP-seq experiment with input controls (left to right) corresponding to ‘unique peaks’ called for each combination of factor as identified in Figure 3F.

**Figure S3 related to Figure 4.**
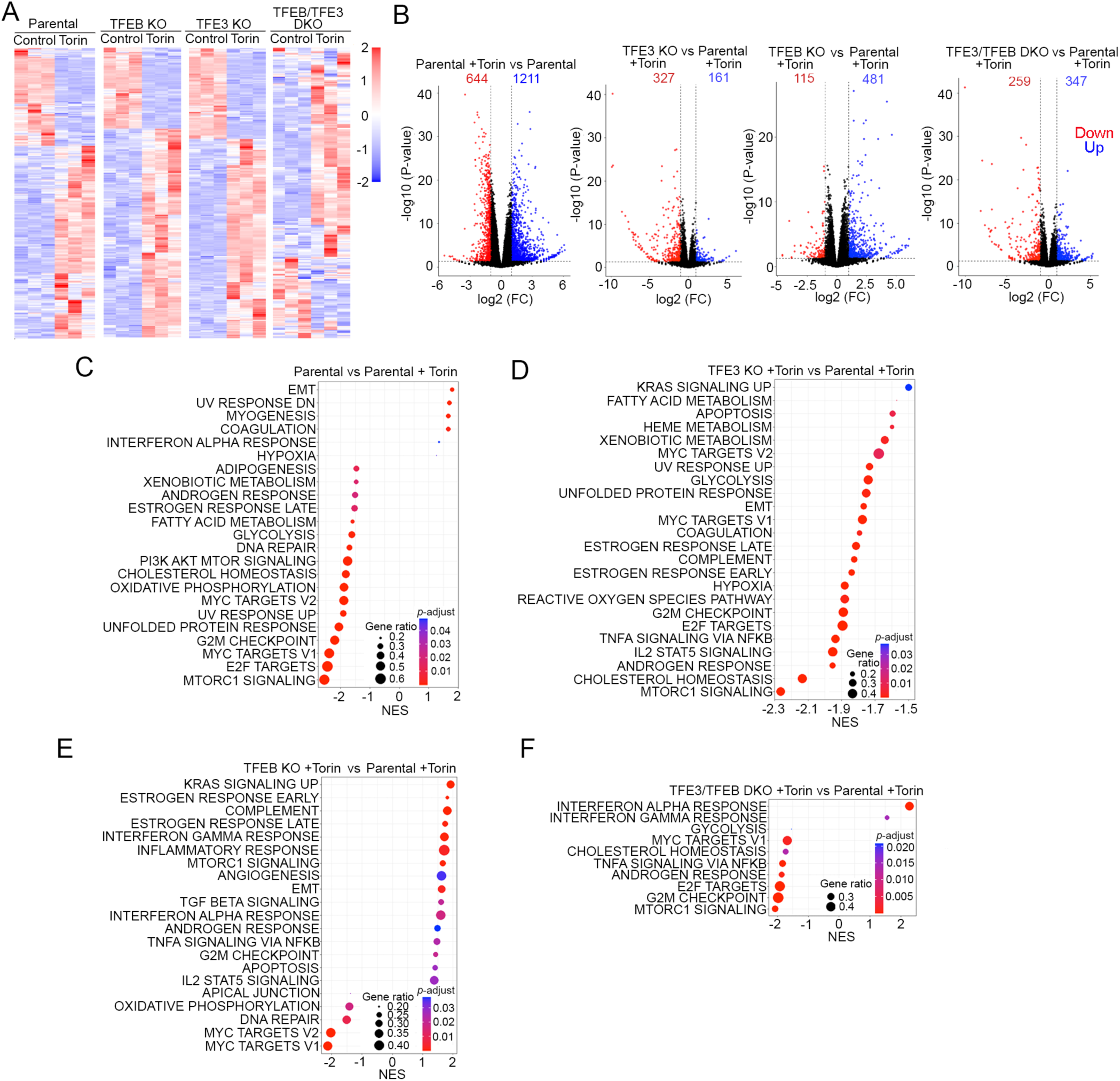
Differential regulation of genes in parental or TFEB/TFE3 KO 501mel cells after Torin treatment. (A) Heatmaps based on triplicate RNA-seq showing differential gene expression of parental or indicated 501mel *TFEB* and *TFE3* KO cell lines before or after treatment with 250 nM Torin for 12 h. (B) Volcano plots showing numbers of significantly (FC ≥ 2, *p* = <0.05) differentially expressed genes comparing parental cells and indicated 501mel KOs in the presence or absence of 250 nM Torin for 12 h as indicated. (C-F) GSEA plots showing significantly differentially enriched gene sets comparing parental or indicated 501mel *TFEB* and *TFE3* KO cells treated or not with 250 nM Torin for 12 h, as indicated.

**Figure S4 related to Figure 5.**
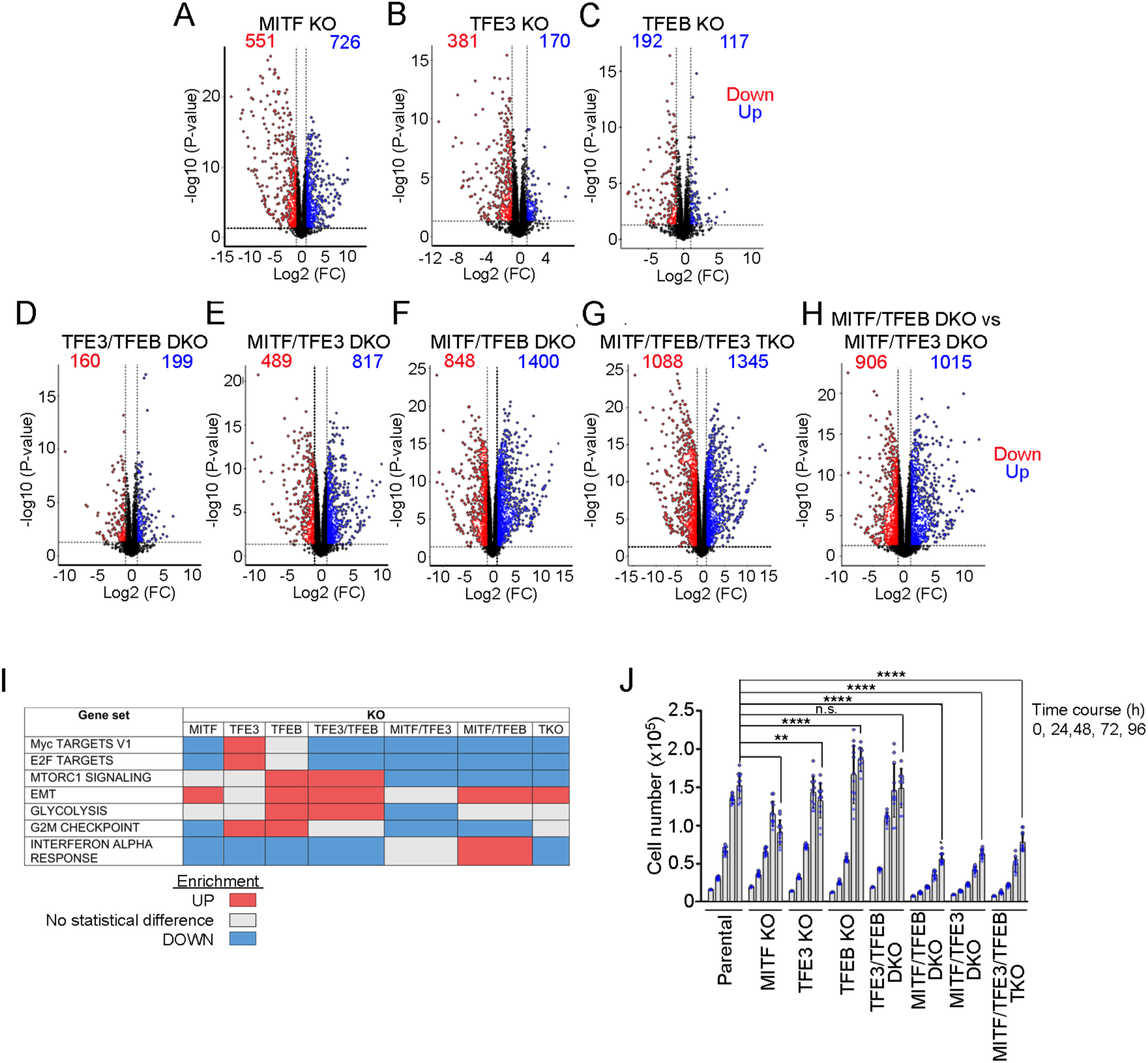
MITF/TFE factors control proliferation. (A-H) Volcano plots showing numbers of significantly (FC 2". 2, *p* = <0.05) differentially expressed genes comparing parental cells and indicated 816-F10 KO cells. (I) Summary of GSEA analyses presented in Figure 5. (J) CyQuant assays of parental or indicated *MITF, TFEB* and *TFE3* B16-F10 KO cells over time after plating of 4,000 cells at time =0 h. N=12 (3 biological replicates in quadruplicate). Error bars indicate S.D. **= *p* < 0.01; **** = *p* < 0.0001; *ns* = not significant when compared to WT cells at 96 h. Linear Mixed-Effects Models was used first to model the slopes then a post-hoc comparison using estimated marginal means accounting for two interacting factors (cell line and time) was used for pairwise comparison. The Bonferroni method was used to adjust p-values for multiple comparisons.

**Figure S5 related to Figure 5.**
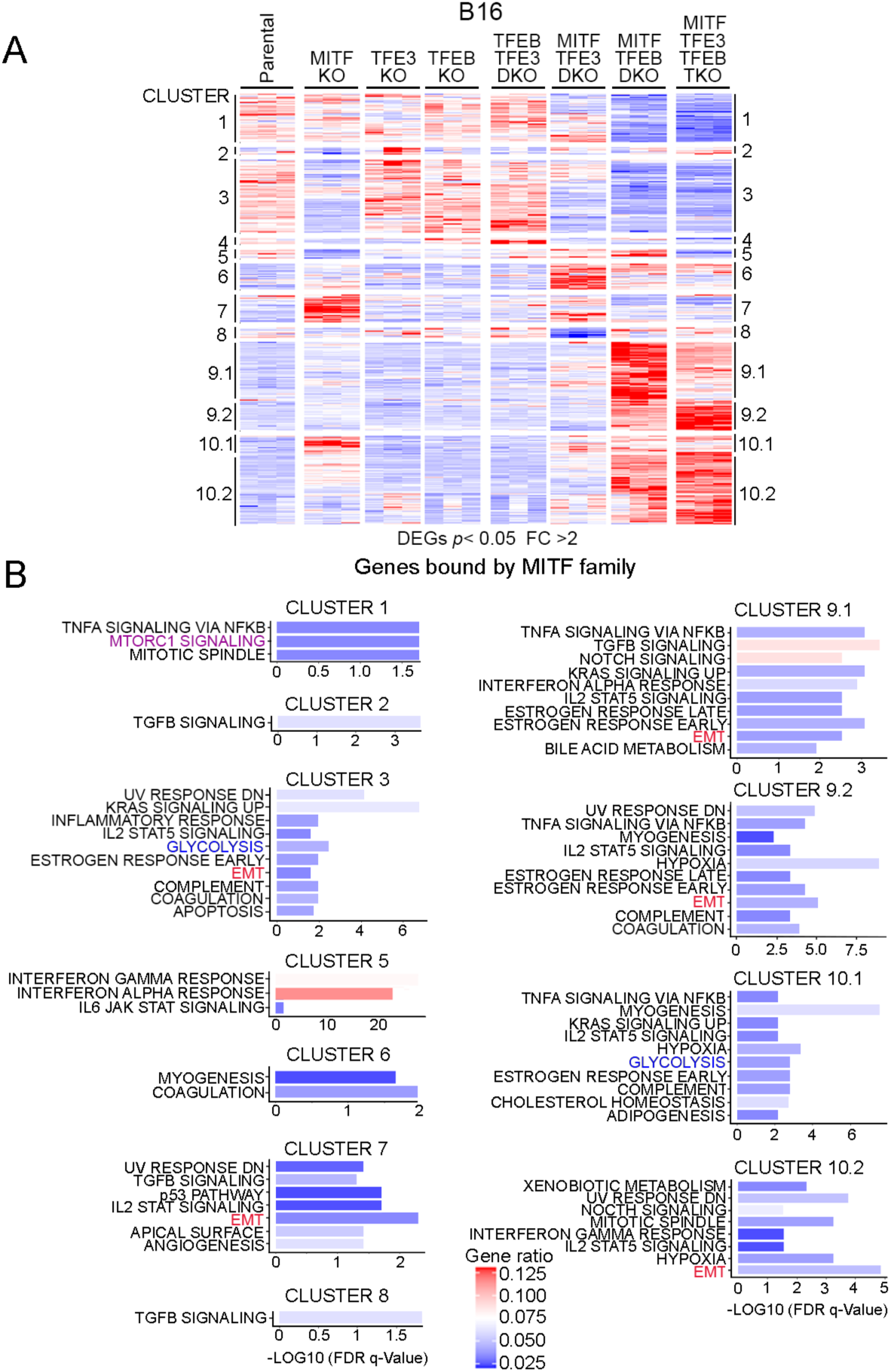
Differential regulation of genes bound by all MITF family members. (A) Heatmaps based on triplicate RNA-seq showing differential gene expression between parental B16-F10 cells and indicated *MITF, TFEB* and *TFE3* KO cell lines. Only those genes commonly bound by all three factors (shown in Figure 3F) are used in the analysis. (B) Enrichment of indicated gene sets within each cluster corresponding to those identified in panel (A) Related to Table S3

**Figure S6 related to Figure 6.**
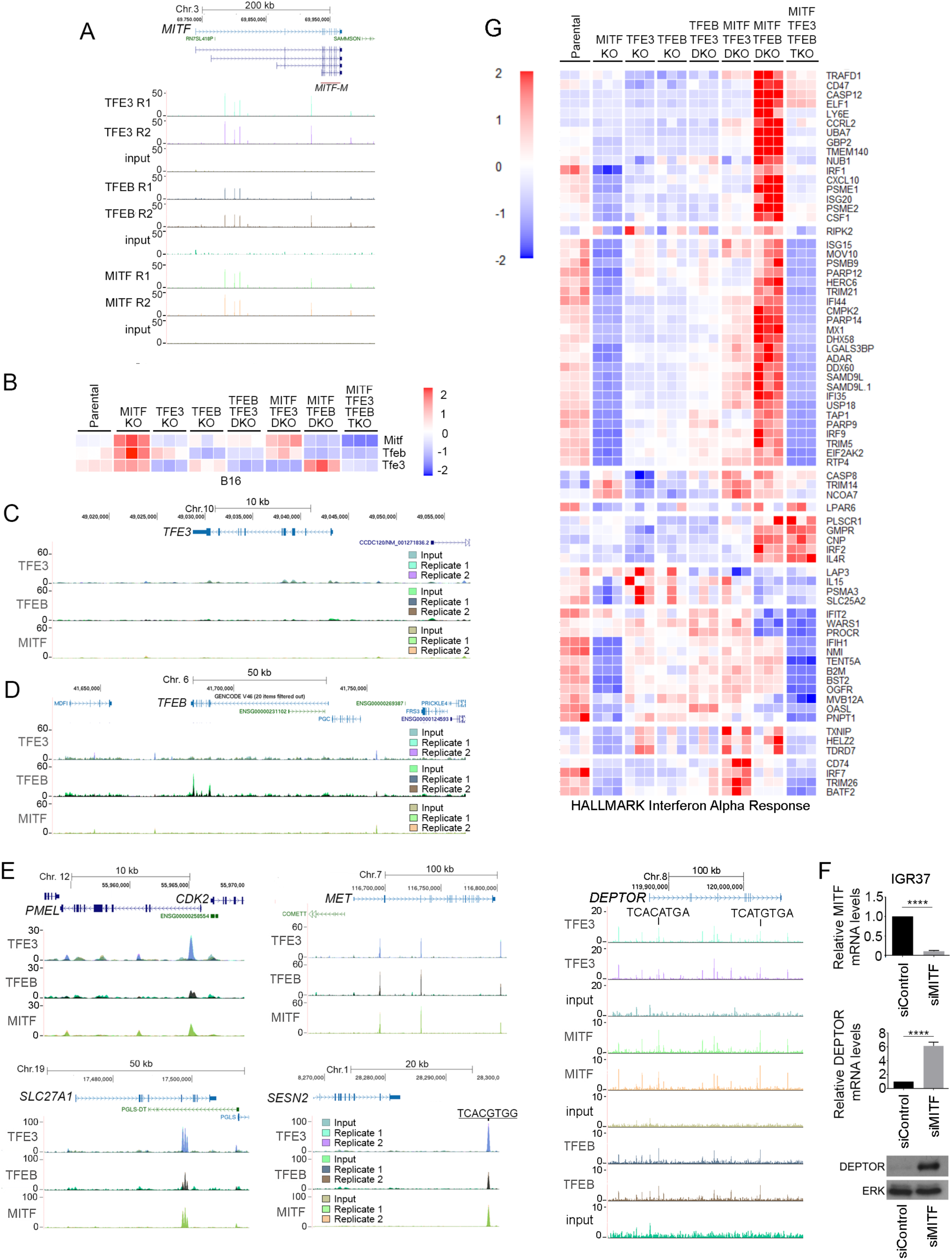
Binding by MITF family members to key genes and regulation of the Interferon Alpha Response gene set. (A) UCSC genome browser screenshots showing binding by MITF, TFEB and TFE3 at the *MITF* locus. (B) Heatmap based on triplicate RNA-seq showing relative expression of *MITF, TFEB* and *TFE3* in indicated parental and B16-F10 KO cell lines. (C-D) UCSC genome browser screenshots showing ChlP-seq profiles of HA-TFE3 and HA-TFEB and HA-MITF at the indicated loci. Binding by TFEB to the *TFEB* gene in panel (D) primarily represents an input signal. (E) UCSC genome browser screenshots showing ChlP-seq profiles of HA-TFE3 and HA-TFEB and HA-MITF at the indicated loci. (F) Results of quantitative RT PCR for *MITF* and *DEPTOR* mRNAs in IGR37 human melanoma cells transfected with control or MITF-specific siRNA. N=3. Error bars = S.D. **** = *p* < 0.0001. Lower panel shows the corresponding western blot. (G) Heatmap showing relative gene expression based on triplicate RNA-seq of parental B16-F10 and derived *MITF, TFEB* and *TFE3* KO cells showing genes from the HALLMARK Interferon Alpha Response gene set. Related to Table S4

